# TBX3 is required for the differentiation of a subset of retinal ganglion and cone photoreceptor cells

**DOI:** 10.64898/2026.10.10.758237

**Authors:** Joao Carlos Ribeiro-Silva, Rachel Burt, Nikita Shah, Mark L Derbyshire, Sruti Akula, Zoe Boyd, Carmen Lee-Bennett, Michael E. Zuber, Andrea S. Viczian

## Abstract

Retinogenesis is a key step driving the formation of the neurons that later coordinate angiogenesis in the retina. Understanding these complex interactions requires dissecting apart its cellular and molecular drivers. To tease apart the molecular links coupling retinogenesis to vascular development, we utilized conditional knockout mouse models to selectively delete the transcription factor T-box 3 (*Tbx3*) in 1) astrocytes, 2) postnatal retinal ganglion cells (RGCs), or 3) retinal progenitors. We demonstrate that the hypovascular retina phenotype observed upon *Tbx3* loss most likely stems from a disruption of early retinogenesis. Embryonic analyses revealed that TBX3 is required during a critical retinogenic window coinciding with ATOH7 activity. While ATOH7^+^ progenitors are more abundant in *Tbx3* mutants, the expression of key RGC and cone precursor factors (including TBX5, RXRγ, ISL1/2, POU4F2, SHH, LHX4 and OTX2) is severely diminished, halting proper RGC and cone differentiation. Utilizing birthdating experiments and developmental immunostaining, we establish TBX3 as a novel upstream regulator of RGC and cone photoreceptor specification. Because the degeneration of RGCs is the definitive hallmark of diseases like glaucoma, identifying TBX3 as a driver of early RGC formation positions it as a promising new candidate biomarker for early-stage glaucoma diagnosis and potentially therapeutic targeting.

**Summary statement:** This study demonstrates for the first time that TBX3 plays a role in mammalian retinogenesis and in the establishment of retinal ganglion cell and cone photoreceptor subtypes.

## INTRODUCTION

Retinal neurons coordinate angiogenesis to build the tri-layered vascular plexus essential for their function and survival. In the superficial layer, retinal ganglion cells (RGCs) recruit astrocytes to guide vascular growth via vascular endothelium growth factor-A (VEGF-A) (Fruttiger et al., 1996; Dorrell et al., 2002; Gerhardt et al., 2003), a process disrupted by RGC loss or altered intrinsically photosensitive RGC (ipRGC) activity (Edwards et al., 2012; Rao et al., 2013). In the deep layer, VEGF gradients from Müller glia (MG) and photoreceptors drive vessel formation (Stone et al., 1995; Rattner et al., 2019), whereas in the intermediate layer, vessel formation is controlled by amacrine and bipolar cell-derived VEGF (Usui et al., 2015). Accordingly, selective ablation of starburst amacrine cells (Weiner et al., 2019) or bipolar cells and photoreceptors (Biswas et al., 2024) leads to severe plexus deficits. Through these layer-specific signaling cascades, retinal neurons actively construct the very vasculature required to sustain them.

We have previously demonstrated that conditional knockout (cKO) of the transcription factor T-box 3 (*Tbx3*) leads to a severe postnatal reduction in dorsal RGCs and ipRGCs, which is paralleled by impaired astrocytic lattice formation and deficient superficial and deep vascular plexus formation (Fig. 1; Derbyshire et al., 2023). Despite this strong correlation, it remains to be elucidated whether the defective angiogenesis in *Tbx3* cKO is a primary consequence of *Tbx3* loss in astrocytes or a consequence of a primary defect in retinogenesis. To address this central question, a deeper understanding of the *Tbx3* role in retinal ganglion cell (RGC) specification is required.

**Figure 1.**
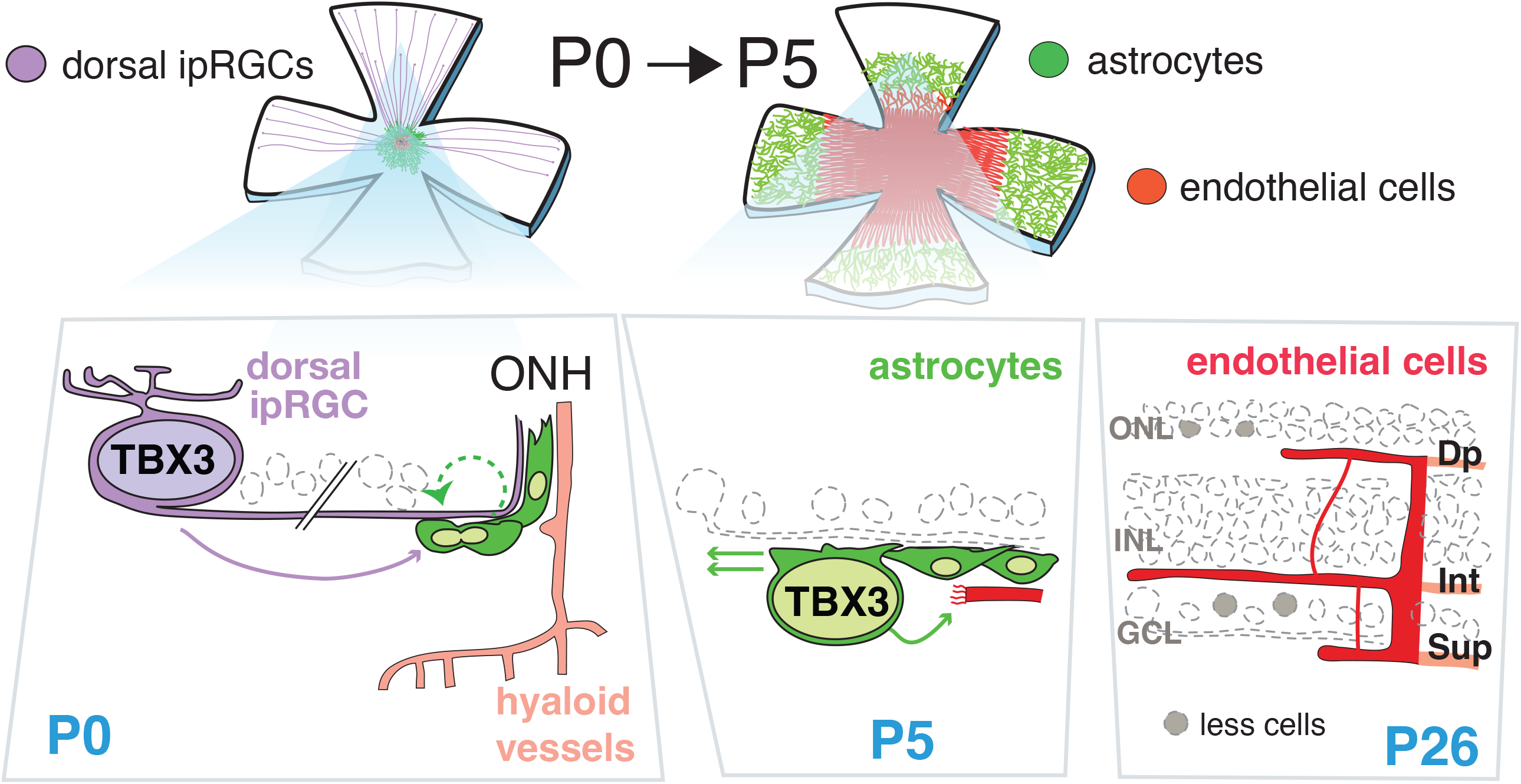
TBX3 is required for dorsal retinal angiogenesis via its activity in dorsal retinal astrocytes and a subset of RGCs. Our previous study showed that TBX3 is required by dorsal ipRGCs, which draw TBX3-expressing astrocytes into the retina, but without TBX3, they fail to populate the dorsal periphery. Astrocytes normally cause sprouting of endothelial cells at the optic nerve head to form the stereotypical tri-layered retinal vasculature (superficial, Sup; deep, Dp; intermediate, Int). Lack of *Tbx3* disrupts this process, causing malformation of both superficial and deep layers in the dorsal retina. Postnatal day 0 (P0), 5 (P5) and 26 (P26).

*Tbx3* is a member of T-box family of transcription factors, characterized by their highly conserved DNA-binding domains. *Tbx3* has established roles in embryonic development, especially as a keeper of pluripotency and as a patterning gene during cardiac, limb, optic and otic development (Suzuki et al., 2004; Yang et al., 2006; Frank et al., 2012; Singh et al., 2012; Kaiser et al., 2021; Derbyshire et al., 2023; Song and Morrow, 2023). In the embryonic eye, *Tbx3* is expressed in the dorsal territory of the optic cup, and it is thought to specify dorsal identity and suppress ventral programs (Sowden et al., 2001; Takabatake et al., 2002; Wong et al., 2002). The timely expression of *Tbx3* in the developing retina at E10.5 (Derbyshire et al., 2023) overlaps precisely with the critical window for the birth of early-born retinal cells, which starts at E11 for RGCs (Brown et al., 1998; Gan et al., 1999) and cone photoreceptors (Carter-Dawson and LaVail, 1979).

In the mouse, the development of first-born retinal neurons begins when naïve retinal progenitor cells (RPCs) acquire the expression of Atonal homolog 7 (ATOH7); Yang et al., 2003; Feng et al., 2010; Brzezinski et al., 2012) and exit the cell cycle (Turner et al., 1990). This process is followed by lineage commitment and cell differentiation. For RGCs, lineage commitment and differentiation rely on the Insulin gene enhancer protein 1 (ISL1) and the POU domain, class 4, transcription factor 2 (Mu et al., 2005; Mu et al., 2008a; Gao et al., 2014; Clark et al., 2019; Wu et al., 2021). As for photoreceptors, this process is mediated by Orthodenticle Homeobox 2 (OTX2), cone rod homeobox (CRX), Prdm1 (BLIMP1) and LIM homeobox protein 4 (Nishida et al., 2003; Katoh et al., 2010a; Brzezinski et al., 2013; Emerson et al., 2013; Buenaventura et al., 2018).

Nonetheless, several lines of evidence demonstrate that our understanding of the mechanisms underlying these phenomena remain incomplete: 1) depletion of ATOH7-dependent mechanisms can be rescued by expression of Bax (Brodie-Kommit et al., 2021); 2) RGCs still form in the absence of *Atoh7*, *Pou4f2*, or *Isl1* (Xiang, 1998; Gan et al., 1999; Wang et al., 2000; Pan et al., 2008; Brodie-Kommit et al., 2021); 3) the RGC transcriptome in knockout models of these genes is partially preserved (Mu et al., 2004; Mu et al., 2005; Li et al., 2014; Wu et al., 2015); and 4) although extrinsic factors, like thyroid hormone and retinoic acid signaling pathways, have been shown to modulate the formation of cone photoreceptor subtypes, the genetic factors that drive their identity remain unknown (Ng et al., 2001; Roberts et al., 2005; Roberts et al., 2006; Applebury et al., 2007).

Since evidence demonstrate the existence of alternative RGC generation pathways (Jiang et al., 2013; Chang et al., 2017; Kuwajima et al., 2017) and *Tbx3* loss results in postnatal loss of dorsal RGCs and ipRGCs (Derbyshire et al., 2023), we hypothesize that TBX3 is a novel regulator of retinogenesis. We further propose that TBX3-dependent mechanisms for neuronal specification might subsequently impair astrocyte migration and the proper establishment of the tri-layered retinal vascular plexus in the mouse.

## METHODS

### Mice

Conditional deletion of *Tbx3* was achieved by crossing the *Tbx3^ΔFl/ΔFl^* transgenic strain (Derbyshire et al., 2023) with the following commercially available mice strains: *Dkk3-Cre* (Tg(Dkk3-cre)^D9Tfur^; RIKEN BioResource Research Center #RBRC05427) (Sato et al., 2007; Derbyshire et al., 2023) and *Six3-Cre* (Furuta et al., 2000) for the optic cup, *Slc17a6-Cre* for mature RGCs (Vong et al., 2011; Sripinun et al., 2025), and *Pdgfrα-CreER^T2^* for astrocytes (Mudhar et al., 1993; O’Rourke et al., 2016; Chung et al., 2018). The specificity of Cre activity was determined by Ai9 cre reporter expression (JAX stock #007909). For each of the generated strains, wild type (Cre^negative^;*Tbx3^Fl/Fl^*) and conditional knockout (Cre^+^;*Tbx3^ΔFl/ΔFl^*) littermates were used, except for the *Dkk3-Cre,* in which sibling wild type (WT) were *Dkk3-Cre^negative^;Tbx3^Fl/+^* and conditional knockouts (cKO) were *Dkk3-Cre*^+^;*Tbx3^ΔFl/ΔFl^*. For the *Pdgfrα-Cre^ER-T2^* experiments, a single tamoxifen injection of 50 μL at 75mg/ml (Sigma Aldrich, #T5648, diluted in corn oil, Sigma Aldrich #C8267) at P0 was used to induce Cre activity. Both WT and cKOs equally received injections. The mice with GFP knocked into the *Tbx3* gene locus (*Tbx3^GR/+^*) were described previously (16). All animals were kept on a C57Bl/6J background. Genotyping was carried out as previously described (Derbyshire et al., 2023). This study was approved by the Upstate Institutional Animal Care and Use Committee and adhered to the ARVO Statement for the Use of Animals in Ophthalmic and Vision Research.

### Tissue isolation and immunostaining

Timed pregnancies were performed by overnight harem breeding and weighing dams, as described (Heyne et al., 2015). Embryos were extracted from the euthanized dam. Embryonic heads were fixed in 4% paraformaldehyde for 1h on ice, subjected to a sucrose series (5%, 10% and 20%) incubating at 4°C overnight, and then mounted in Tissue-Tek O.C.T. Compound (Sakura Finetek. Torrance, CA, USA) and cryosectioned (12 μm). The sections were immunostained overnight with the antibodies indicated in Table 1 and covered with glass coverslips with FluorSave (Sigma-Millipore) and 2% 1,4-Diazabicyclo[2.2.2]octane (DABCO; Sigma-Aldrich). For antigen retrieval, cryostat sections were rehydrated in 1X PBS and then boiled in 10mM Sodium Citrate plus 0.05% Tween-20, pH 6.0 for 15 minutes and cooled for 30 minutes at room temperature (RT) before adding blocking solution and proceeding with antibody staining, as previously described (Derbyshire et al., 2023). Birthdating of RGCs was performed by injecting 5-ethynyl-2’-deoxyuridine (EdU, ChemCruz, sc-284628A, 12μg/g body weight, diluted in 1x PBS) in pregnant dams at E12.5, followed by tissue collection and EdU detection using the Invitrogen™ Click-iT™ EdU Cell Proliferation Kit for Imaging, Alexa Fluor™ 488 dye (Invitrogen #C10323) and standard immunostaining analysis, as described in (Miesfeld et al., 2020). Retinal flat mounts were collected and stained with the indicated antibodies, placed within reinforcement labels (Avery Dennison #5722) on glass slides and covered with FluorSave and 2% DABCO and round glass coverslips.

### *Ex-vivo* retina culture

*Ex-vivo* retina culture was performed as described in (Kaufman et al., 2019). Briefly, timed pregnancies were used to obtain sibling WT and cKO E12.5 embryos. The eyes were extracted and cultured in HBSS with calcium, magnesium, HEPES and glucose (Sigma Aldrich #55037C) for 24h under soft agitation in neurobasal media (Gibco™ #21103049) supplemented with 1% N2, 1% fetal bovine serum (FBS; Gibco™ #A5256701), 1% penicillin/streptomycin (Corning #30-001-CI), 1% glutamax (Gibco™ #35050–061) and 10 μM of γ-secretase inhibitor IX (Calbiochem, #565770) diluted in DMSO. Ninety minutes before the 24h-period, 20ng of EdU diluted in 1x PBS (Biosynth Carbosynth #NE08701) was added to the media. After this period, eye cups were fixed in 2% PFA for 20min, subjected to a sucrose series (10%, 20% and 30%), mounted in Tissue-Tek O.C.T. Compound (Sakura Finetek. Torrance, CA, USA), cryosectioned (12 μm) and stained according to the conditions indicated in Table 1.

### Primary astrocyte culture and western blot analysis

Primary P6 retinal astrocytes were isolated as previously described with modifications (Kirschner et al., 2021; Strat et al., 2025). Briefly, the optic nerve heads were dissected in sterile PBS and transferred to ice-cold complete DMEM/F12 media (Invitrogen 113330-032), supplemented with 10% FBS, 1% Pen/Strep, 1% Glutamax, and 25 ng/mL EGF (Sigma #E4127– 5X). Tissue underwent mild enzymatic digestion with 0.05% Trypsin/EDTA (Sigma Aldrich #25200-56) for 10 min at 37°C. Trypsinization was halted by adding culture media, and the suspension was centrifuged at 1000 rpm for 5 minutes to pellet the tissue. Digested tissue was resuspended and plated onto 12-well plates pre-coated with 0.2% gelatin (Sigma Aldrich #G1393). Cells were cultured for one week until confluent, then passaged onto 100 mm dishes. To maximize protein yield, a pool of WT and cKO astrocytes was used for the final lysis step. Cells were lysed on ice using RIPA lysis buffer supplemented with protease and phosphatase inhibitors. Homogenates were cleared via centrifugation at 15,000 x *g* for 10 minutes at 4°C. Total protein concentration was determined using the Pierce Rapid Gold BCA Protein Assay Kit (ThermoFisher #A53225). Protein was separated by SDS-PAGE using a 12% resolving gel run for 1 hour at 120 V, then wet-transferred to a PVDF membrane and blocked under agitation for 1h at room temperature. The membrane was incubated with rabbit anti-TBX3 (Invitrogen, #42-800 1:1000 v/v) and mouse anti-β-actin (Li-Cor, #926-4212 1:1000 v/v) antibodies overnight at 4°C with gentle agitation, followed by 1-hour incubation with appropriate secondary antibodies at room temperature. Bands were visualized using the Odyssey CLx Infrared Imaging System (Li-Cor).

### Imaging

Images were obtained in a Leica DM600B fluorescence microscope (Leica, Wetzlar, Germany) coupled to a MicroPublisher 6 camera (Teledyne Photometrics, Tucson, AZ, USA). Retina flatmount images were stitched with the Volocity v6.3 software. Whenever possible, WT and cKO samples were placed on the same slide and images were taken using the same exposure conditions and equally processed using Photoshop (Adobe, San Jose, CA, USA) for brightness and contrast. Cells were counted by individuals blind to the mouse’s genotype. Vascular morphometric parameters were obtained with AngioTool (Zudaire et al., 2011).

### Bulk RNA-Seq

Total RNA was extracted using the miRNeasy Micro Kit (Qiagen #217084) from E13.5 retinas isolated from two litters of sibling wild type (n=3) and cKO (n=3) embryos. Samples were sent to Admera Health for processing. Isolated RNA sample quality was assessed by High Sensitivity RNA Tapestation (Agilent Technologies Inc., California, USA) and quantified by Infinite F Nano+ 200 Pro Tecan (Tecan, Switzerland). All samples were prepared using NEBNext® Poly(A) mRNA Magnetic Isolation Module and NEBNext® Ultra™ II Directional RNA Library Prep Kit for Illumina® (New England BioLabs Inc., Massachusetts, USA). Final libraries quantity was assessed by Qubit 2.0 (ThermoFisher, Massachusetts, USA) and quality was assessed by High Sensitivity D1000 TapeStation ScreenTape (Agilent Technologies Inc., California, USA). Final library size was between 430 bp to 460 with an insert size of 300 to 330bp. Illumina® 8-nucleotide dual-indices were used. Equimolar pooling of libraries was performed based on QC values and sequenced on an Illumina® NovaSeq X plus (Illumina, California, USA) with a read length configuration of 150 PE for 40 M PE reads per sample (20M in each direction). RNA-seq data were analyzed using Partek^TM^ Flow^TM^ software, v12.10. Adapters were trimmed from raw reads using CUTADAPT v4.2. Base trimming from both ends was performed based on quality score (Phred score) of 35 or greater, with a minimum read length of 25. Reads were aligned to *Mus musculus* ver. mm39 using STAR v2.7.8 aligner, and counts were generated using Ensembl release v115. Features with a count sum ≤ 3.0 were excluded. Differential expression analysis was performed using DESeq2. P values were adjusted using the Benjamini-Hochberg method, and genes with FDR ≤ 0.05 and a fold change ≥ 2.0 were considered differentially expressed. Heatmaps were generated from normalized expression values in R using the pheatmap package, version 1.0.13 (Kolde R. *pheatmap: Pretty Heatmaps*. R package version 1.0.13. 2025. Available from: https://github.com/raivokolde/pheatmap).

### Statistical Analysis

Data represented as mean ± standard error of the mean (SEM). Symbols represent the average number of the density of cells counted from both eyes per mouse (n), where counts were taken from at least three sections from each eye. Volume measurements were taken with Volocity v6.3 (Perkin Elmer). Data was normalized by the average value of the sibling wild type in each litter. Graphpad Prism® v10 was used for statistical analysis. Cell counts and volumes were measured under double blind conditions. Sample size was determined by an a priori power analysis in G*Power (G*Power 3.1) using a two-tailed independent-samples *t*-test (alpha = 0.05, power = 0.80). The effect size was estimated in G*Power using previously published data.

## RESULTS

### TBX3 is required in RPCs, not in mature RGCs or astrocytes, to ensure retinogenesis and angiogenesis

Loss of *Tbx3* at the optic cup stage impairs ipRGC formation, astrocyte migration, and dorsal retinal angiogenesis (Fig. 1; Derbyshire et al., 2023). However, because *Dkk3-Cre* targets both neural progenitors and astrocytes, the underlying cellular mechanisms remain unclear. We hypothesized that these defects stem from a loss of *Tbx3* in RPCs rather than in differentiated RGCs or astrocytes. To test this, we used conditional deletion models to systematically isolate the contributions of each cell lineage to the *Tbx3* cKO neurovascular phenotype.

We first conditionally deleted *Tbx3* in differentiated RGCs (*Tbx3^ΔRGCs^*) using the *Slc17a6-Cre* (Tran et al., 2019; Sripinun et al., 2025) whose activity was detected as early as E18.5 (Fig. S1A-B’), in agreement with the perinatal onset of endogenous *Slc17a6* expression (Johnson et al., 2007). At P9, *Tbx3^ΔRGCs^* retinas showed no overt defects in the astrocytic lattice and vascular morphometric measurements (Fig. 2A-C; Fig. S1C-H). We also observed no difference in RGCs density when comparing *Tbx3^ΔRGCs^* to sibling WT mice with ISL1/2 (Fig. 2D-H) or BRN3A (Fig. S1I-M) staining. These findings demonstrate that *Tbx3* expression by differentiated RGCs is dispensable for RGC density, astrocyte migration, and retinal angiogenesis.

**Figure 2.**
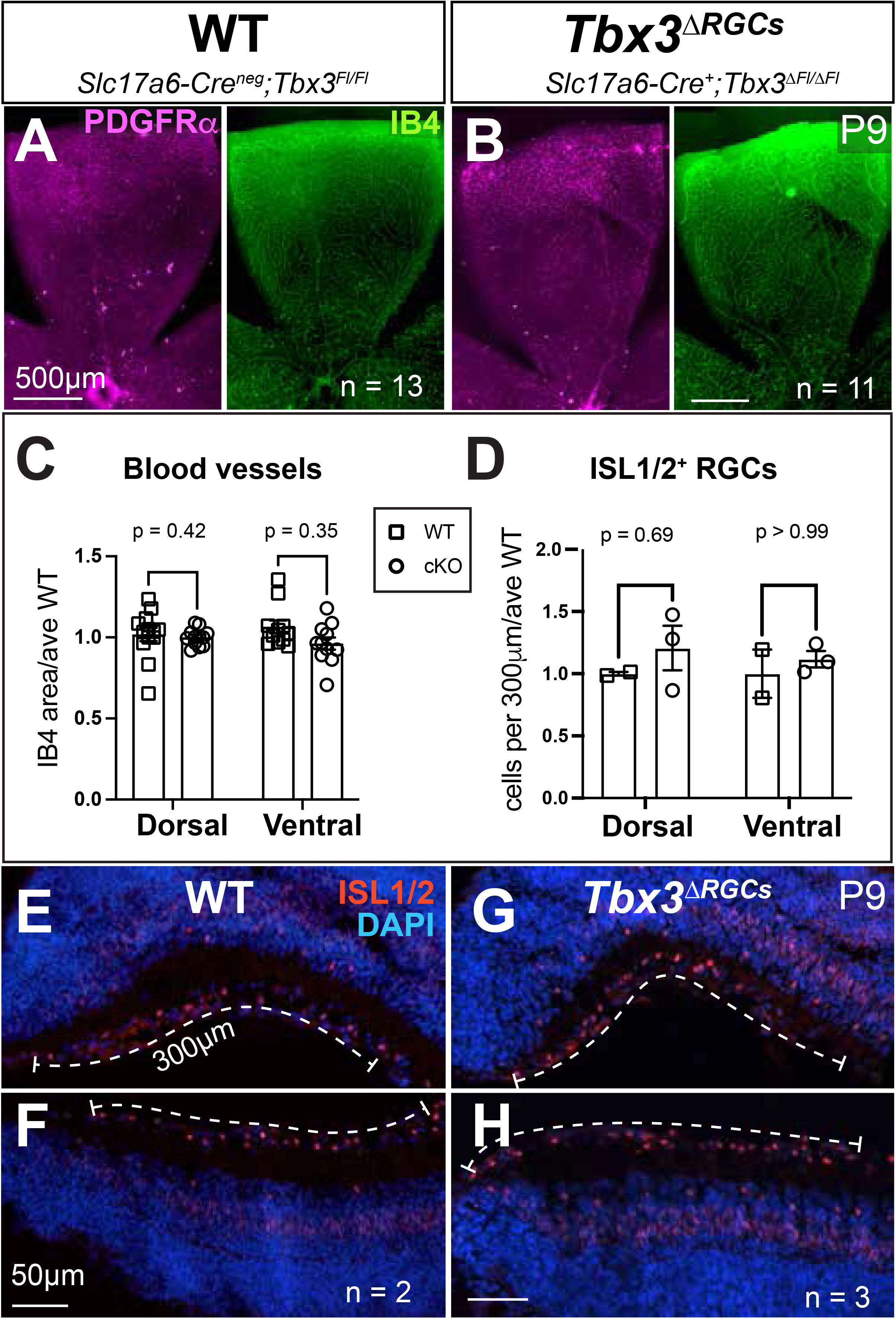
TBX3 is not required by differentiated RGCs for retinal angiogenesis. (A, B) P9 dorsal retinal flat mounts from WT (A) and *Tbx3^ΔRGCs^* (B) littermates stained for PDGFRα (astrocytes, magenta) and IB4 (vessels, green). (C) Retinal % vessel area quantification. (D) Quantitation of ISL1/2^+^ RGCs within 300 µm dorsal and ventral edges of central retinal sections. (E–H) P9 central retinal sections from WT (dorsal, E; ventral, F) and *Tbx3^ΔRGCs^* (dorsal, G; ventral, H) mice stained for ISL1/2 (red) and DAPI (blue). Dashed lines mark the 300 µm measurement regions quantified in (D). Data analyzed by two-way ANOVA with Bonferroni test. Scale bars: 500 µm (A, B), 50 µm (E–H).

Next, we assessed the role of *Tbx3* in astrocytes, which are critical for vascular guidance and that we and others have previously demonstrated to express *Tbx3* (Derbyshire et al., 2023; Clarke et al., 2018). We took advantage of the *PDGFRα-CreER^T2^* (Duan et al., 2014; O’Rourke et al., 2016) to target *Tbx3* as the astrocytes migrate into the retina at P0 (Fig. 3A), thereby preceding the peak formation of MG (Young, 1985). We confirmed Cre activity in immature and differentiated astrocytes (Fig. S2A’-C”) and verified *Tbx3* loss in isolated cultured astrocytes (Fig. S2D). In the absence of *Tbx3*, astrocyte migration is delayed as early as P3 (WT: 1.0 ± 0.015; cKO: 0.67 ± 0.019; p ≤ 0.001) and persisting at P5 (WT: 1.0 ± 0.17; cKO: 0.826 ± 0.015; p ≤ 0.001; Fig. 3B-E, H).

**Figure 3.**
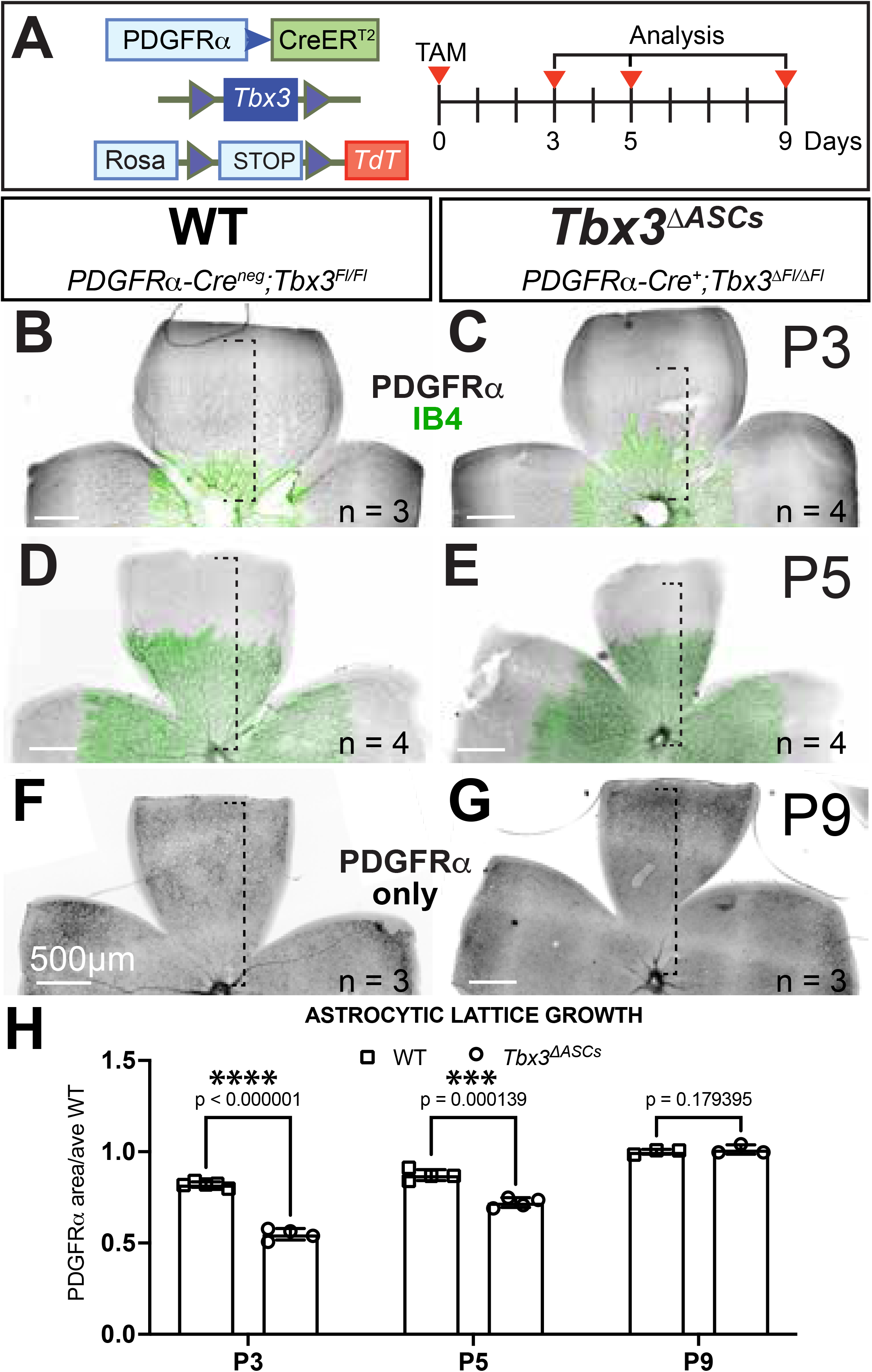
Astrocyte *Tbx3* is not required for retinal angiogenesis. (A) Experimental timeline showing tamoxifen (TAM) induction at P0 and tissue collection at P3, P5, and P9. (B–E) P3 and P5 retinal flat mounts from WT (B, D) and *Tbx3^ΔASCs^* (C, E) stained for astrocytes (PDGFRα, black) and blood vessels (IB4, green). Dashed line: astrocytic area measured. (F, G) P9 retinal flat mounts from WT (F) and *Tbx3^ΔASCs^* (G) mice. (H) Quantification of astrocyte migration distance across time points. Data analyzed by unpaired Student’s t-test; **p = 0.0003, ***p = 0.0001.

However, this delay did not impact angiogenesis, as it progressed similarly between WT and cKO (Fig. S2E). In addition, by P9 *Tbx3^ΔASCs^* astrocytes overcame their migration defect (Fig. 3F-H) and the superficial vascular plexus was indistinguishable from WT (Fig. S2F). These findings indicate that astrocyte *Tbx3* is required for their initial migration in the retina but deny its requirement for the dorsal defects in astrocytic lattice and angiogenesis of the optic cup *Tbx3* cKO.

Finally, we generated *Tbx3^ΔProgen^* mice by targeting optic vesicle progenitors at E9.0 with the *Six3-Cre* (Furuta et al., 2000). At P9, *Tbx3^ΔProgen^* retinas recapitulated the severe *Dkk3-Cre* cKO phenotype (Derbyshire et al., 2023), with a complete absence of the dorsal astrocytic lattice (PDGFRα) and a statistically significant reduction in vascular coverage in the *Tbx3^ΔProgen^* compared to WT (Fig. 4A-B’,G; Fig. S3A,B; WT 90.38 ± 0.94; cKO 54.45 ± 2.5). Interestingly, this mutant reduced dorsal and ventral RGC density (Fig. 4C-F, H; ISL1/2 dorsal: WT 1.0 ± 0.02, *Tbx3^ΔProgen^* 0.49 ± 0.04; ventral: WT 1.0 ± 0.03, *Tbx3^ΔProg^* 0.37 ± 0.09; Fig. S4A-C; BRN3A dorsal: 1.0± 0.01, *Tbx3^ΔProgen^* 0.373 ± 0.06; ventral: WT 1.0 ± 0.02, *Tbx3^ΔProgen^* 0.43 ± 0.12), unlike the dorsal-restricted effect that we observed with the *Dkk3-*Cre (Derbyshire et al., 2023). However, we found mosaic reduction of TBX3 in the embryonic retina of these mice (Fig. S3C-D’), as expected (Furuta et al., 2000), unlike the complete ablation of retinal TBX3 using our previously characterized *Dkk3-Cre* cKO (Fig. S3E-F’; Derbyshire et al., 2023).

**Figure 4.**
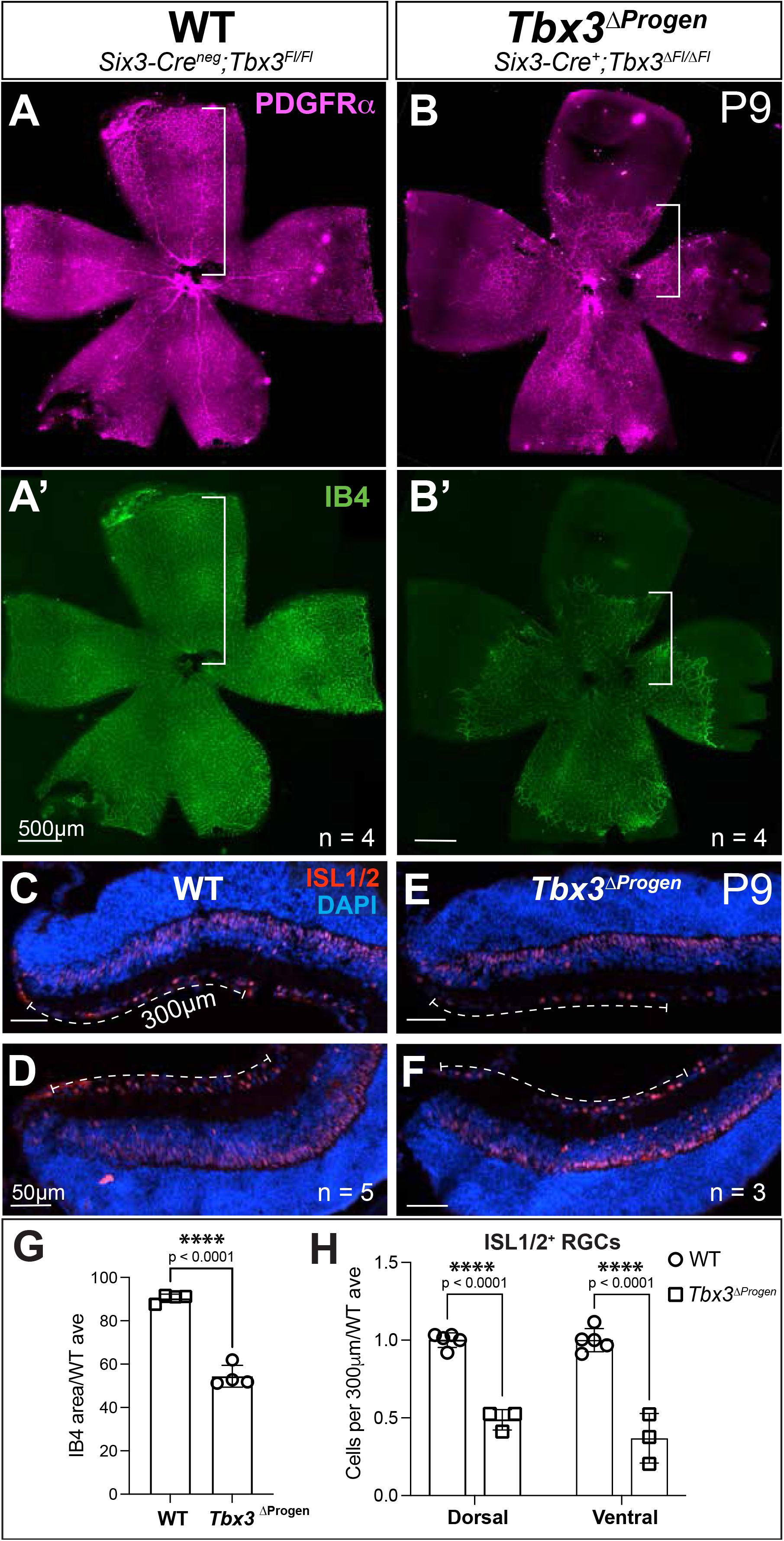
A second Cre-driver targeting astrocytes and RPCs confirms the hypovascular phenotype of *Tbx3* loss. (A, B) P9 retinal flat mounts from WT (A, A’) and *Tbx3^ΔProgen^* (B, B’) mice stained for PDGFRα (astrocytes, magenta; A, B) and IB4 (vessels, green; A’, B’). White bar: vascular migration distance from optic nerve head. (C–F) P9 retinal sections from WT (dorsal, C; ventral, D) and Tbx3*^ΔProgen^* (dorsal, E; ventral, F) mice stained for ISL1/2 (red). Dashed lines mark 300 µm quantification regions. (G) Quantification of vascular coverage. (H) Quantification of ISL1/2^+^ RGCs in dorsal and ventral regions. Data analyzed by unpaired Student’s t-test (G), two-way ANOVA with Bonferroni test (H). Scale bars: 500 µm (A, B), 50 µm (C–F).

Taken together, these findings support the hypothesis that early RPC and not RGC or astrocyte TBX3 may drive the angiogenic phenotype observed at P9. Given the mosaicism of the *Six3-Cre* mouse lineage and the fact that we have thoroughly characterized the *Dkk3-Cre* cKO (Derbyshire et al., 2023), we went back to this model to determine the influence of RPC TBX3 on retinogenesis.

### *Tbx3* loss reduces retinogenesis, increases proliferation and apoptosis

Based on timely expression of *Tbx3* during retinogenesis (as early as E9.0, Fig. S5A), our genetic complementation results and our previous evidence of ipRGC loss upon *Tbx3* cKO, we hypothesized that *Tbx3* loss impairs retinogenesis. To test this, we analyzed retinal differentiation in WT (*Dkk3-Cre^negative^;Tbx3^Fl/+^)* and cKO (cKO: *Dkk3-Cre^+^;Tbx3^ΔFl/ΔFl^*) at E13.5, a stage coinciding with the plateau of ipRGC genesis (Lucas and Schmidt, 2019). Using Class III β-tubulin, an early neural marker (Memberg and Hall, 1995), we observed that while WT retinas showed robust β-tubulin that spanned the innermost developing cell layer (DCL), this pattern was significantly reduced in cKO retinas (WT: 1.0±0.05; cKO: 0.74±0.09; p=0.0167; Fig. 5A-C).

**Figure 5.**
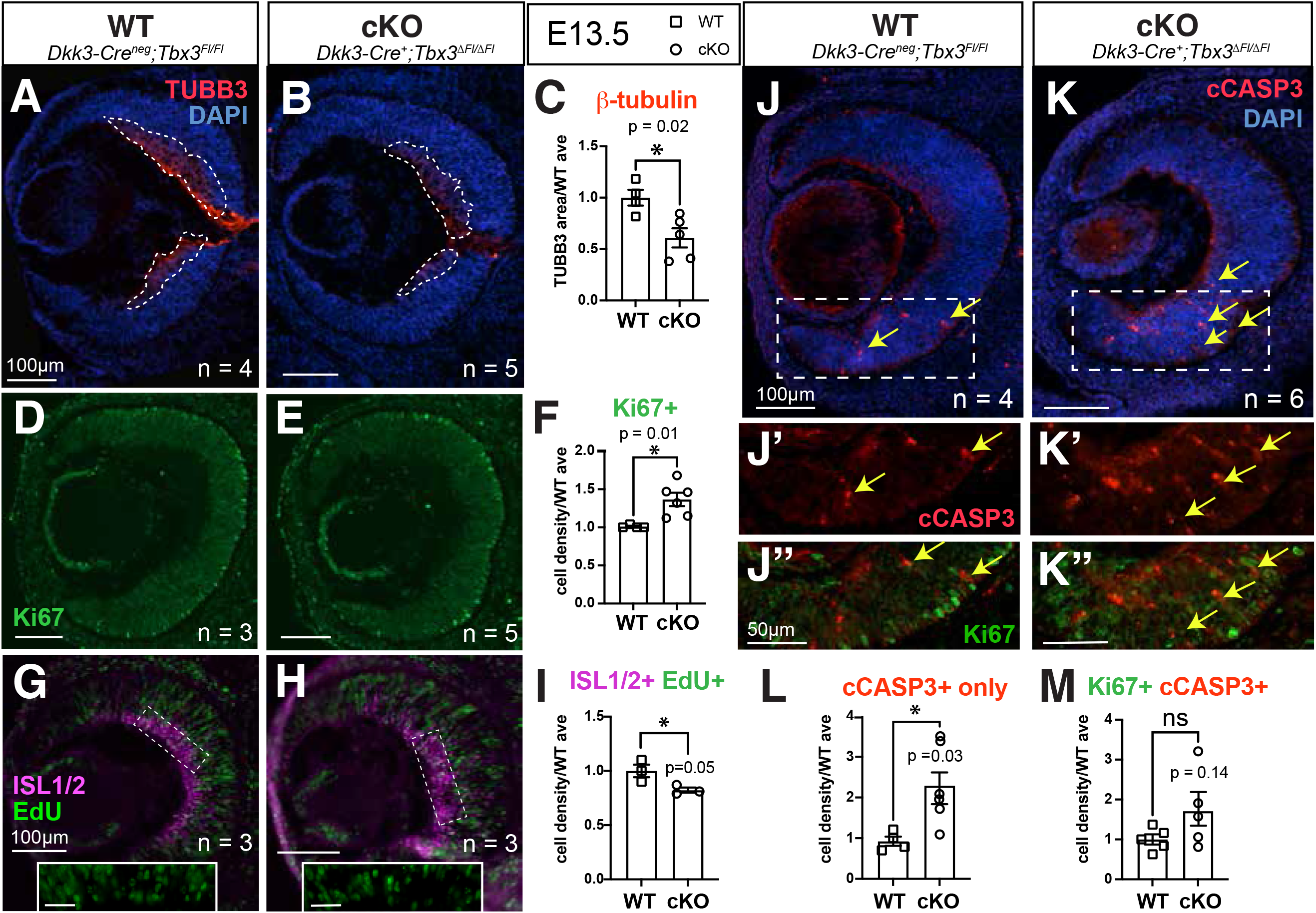
TBX3 regulates neuronal differentiation. (A–C) E13.5 WT (A) and cKO (B) retinal sections stained for TUBB3 (red) with quantification (C). (D–F) E13.5 WT (D) and cKO (E) sections stained for Ki67 (green) with quantification (F). (G–I) RGC birthdating (EdU pulse at E12.5, analysis at E13.5) in WT (G) and cKO (H) retinas with EdU^+^ISL1/2^+^ quantification (I). Insets: boxed areas (scale bar: 25 µm). (J–K″) E13.5 WT (J–J″) and cKO (K–K″) retinas stained for Ki67 (green), cCasp3 (red; yellow arrows indicate positive cells), and DAPI (blue). (L) Quantification of cCasp3^+^ and Ki67^+^cCasp3^+^ cells. Data analyzed by Unpaired Student’s t-test (C, F, I, L). Scale bars are indicated in each panel.

As retinogenesis is limited by RPC proliferation, we quantified cycling RPCs in E13.5 retinas using Ki67 staining. The loss of *Tbx3* caused an ∼40% increase in Ki67^+^ RPCs when compared to WT (WT: 1.0 ± 0.01; cKO: 1.37 ± 0.09; p = 0.012; Fig. 5D-F’”). To gain insight into RPC output, we labeled RPCs with EdU at E12.5 and determined the number of RGC precursors at E13.5 by co-staining EdU with an anti-ISL1/2 antibody. In agreement with the reduced β-tubulin staining, we found decreased RGC precursors in *Tbx3* cKO retinas (EdU^+^ISL1/2^+^ WT: 1.0 ± 0.02; cKO: 0.87 ±0.02; p = 0.048; Fig. 5G-I). To determine if dysregulated RPC proliferation had a causative role in the reduced retinogenesis of the cKO, we cultured E12.5 eye cups in the presence of a γ-secretase inhibitor, limiting Notch signaling and forcing RPCs out of the cell cycle (Nelson et al., 2007). If *Tbx3* regulates the cell cycle exit, this strategy would either increase or equate the number of RGC precursors in the cKO compared to the WT after 24h of treatment (∼ E13.5). The inhibitor’s efficacy was confirmed by the absence of EdU^+^ RPCs in the retina (arrows, Fig. S5B-C); however, it had no impact in the cKO RPC output, which remained poor in RGC precursors when compared to the WT (Fig. S5B’-E).

The uncoupling between retinogenesis and RGC differentiation was associated with increased apoptosis, as evidenced by the increased cleaved-caspase 3 (cCasp3) staining observed in cKO retinas at E13.5 (Fig. 5J-L; WT:0.92 ± 0.11; cKO: 2.29 ± 0.39, p = 0.03). We also observed an overlap between the cCasp3 and Ki67 staining that was trending toward an increase in the cKO (Fig. 5K’-K”,M; WT: 1.0 ±0.13; cKO: 1.7 ±0.42; p = 0.14) but not statistically significant, possibly due to the low number of apoptotic counts at this stage. Collectively, these findings support the hypothesis that *Tbx3* is required for early retinogenesis. Without it, multipotent RPCs persist in the cell cycle, reducing lineage-restricted RGC precursors, and ultimately leading to fewer RGCs in the postnatal retina.

### RNA-sequencing analysis reveals changes in early RGC and cone pathways

To gain insight into the role of TBX3 in retinogenesis, we performed bulk RNA-sequencing (RNA-seq) on WT and cKO E13.5 retinas (Fig. 6). Principal component analysis (PCA) of gene expression demonstrated clustering of samples according to genotype, alongside the expected loss of *Tbx3* expression in cKO samples (Fig. S6A,B). We identified a total of 490 differentially expressed genes (DEGs) between cKO and WT retinas; among these, 201 were downregulated and 289 were upregulated (log2FC>0.55 or <−0.55, q<0.05; Fig. 6A and Dataset S1). Ingenuity Pathway Analysis (IPA) revealed an enrichment of pathways associated with neuroinflammation, macrophage activation, and phagosome formation (Fig. 6B, Fig. S6C), consistent with an increased number of Iba1^+^ cells observed in the cKO retina (Fig. 6C; WT: 1.0±0.02; cKO: 2.1±0.38; p=0.03). Conversely, we observed downregulation of genes associated with stem cell pluripotency, neurogenesis, WNT/SHH signaling, axonal guidance, and synaptogenesis (Supplemental Table S2 and Fig. S6C). While pathways regulating cell proliferation remained unchanged, the pro-apoptotic genes *Casp1* and Bcl2 modifying factor (*Bmf*) were significantly upregulated (Fig. S6D; 5.83-fold and 3.76-fold, respectively; p≤0.007 for both), consistent with the elevated apoptosis observed in E13.5 cKO retinas (Fig. 5K). Together, these results are consistent with a model in which the increased apoptosis trigger microglia activation in E13.5 cKO retinas.

**Figure 6.**
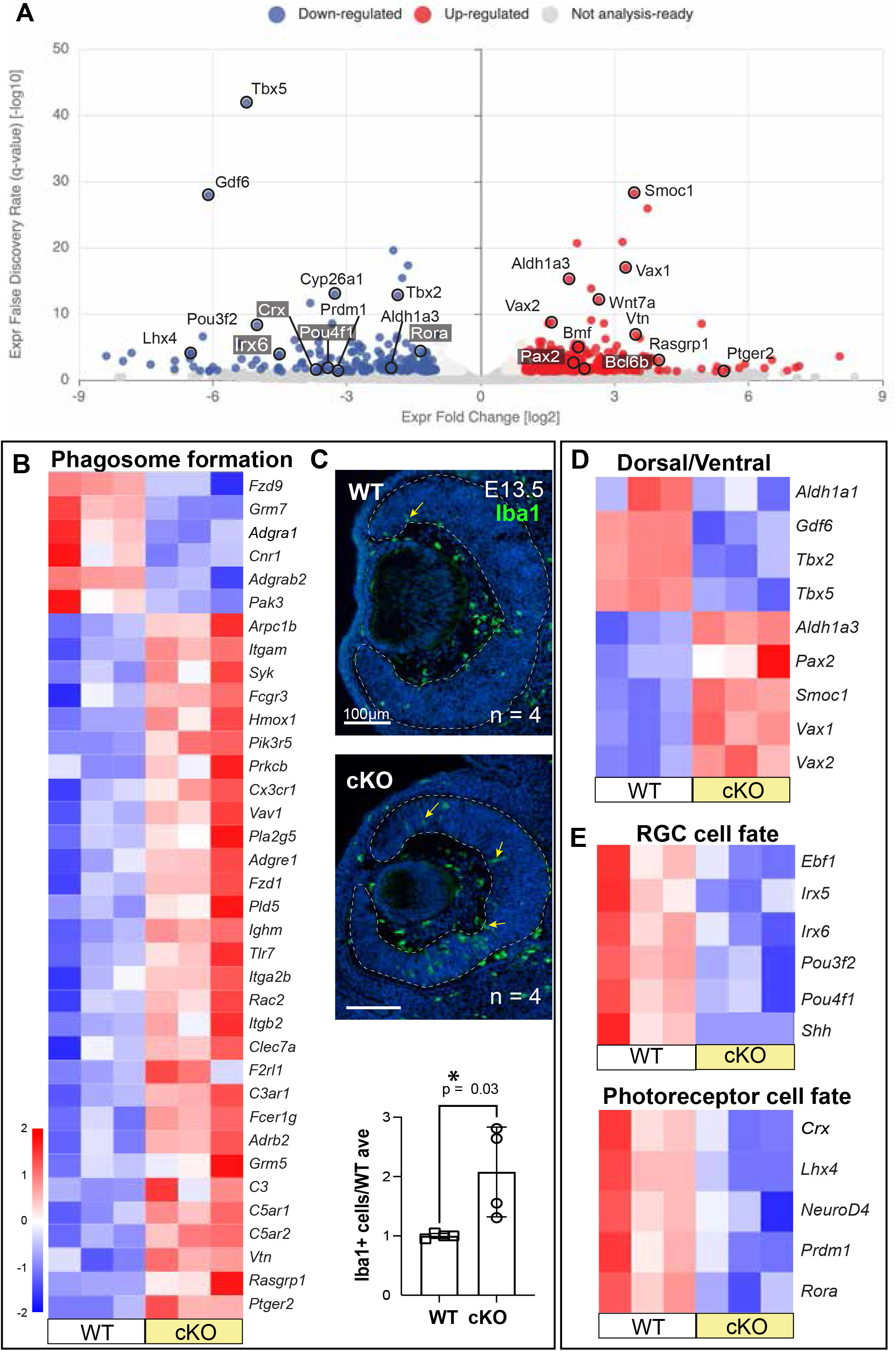
TBX3 is required for expression of RGCs and early photoreceptor genes, and its loss triggers an immune response in the embryonic retina. (A) Volcano plot of differentially expressed genes (DEGs) between E13.5 cKO and WT (n = 3) retinas. (B, C) Heat map of phagocytosis and immune response genes (B) and IBA1^+^ microglia/macrophage immunostaining (C). (D) Heat map of dorsal and ventral retinas DEGs. (E) Heat map of downregulated RGC and photoreceptor genes. Data analyzed by Unpaired two-tailed Student’s t-test (C). Scale bar: 100 µm (C).

Previous studies have established a role for *Tbx3* in patterning the chick and frog eye (Takabatake et al., 2000; Peters and Cepko, 2002; Takabatake et al., 2002; Wang et al., 2015; Ayten et al., 2026). In support of this role in mice, *Tbx3* cKO retinas displayed decreased expression of the dorsal genes *Aldh1a1, Gdf6*, *Tbx2* and *Tbx5,* whereas the ventral markers *Aldh1a3, Pax2, Smoc1, Vax1,* and *Vax2* were upregulated (Fig. 6D). Furthermore, *Tbx3* loss disrupted key signaling pathways driving eye development. We observed a significant reduction in components of BMP (*Bambi, Gdf6, Smad9*), FGF (*Etv1, Dusp4, Fgf8*), Notch (downregulated: *Hey1, Id1, Id2, Id4, Mamld1*; upregulated negative regulator, *Shisa2*) and Shh (*Shh, Ptchd1*) signaling, alongside Wnt pathway genes (up: *Fzd1, Wnt7b;* down: *Fzd9*; Fig. S6D).

Consistent with our findings of reduced retinogenesis at E13.5 and diminished RGCs at P9, *Tbx3* loss correlated with the downregulation of genes downstream the ATOH7-ISL1-POU4F2 RGC differentiation, including *Ebf1, Irx5, Irx6, Pou3f2 (Brn2), Pou4f1 (Brn3a) and Shh* (Wu et al., 2021). These transcriptional changes aligned with the downregulation of genes involved in axon pathfinding, neuronal maturation and synaptogenesis (Fig. S6E). Similarly, we noted a reduction in genes critical for photoreceptor genesis, matching the timeline for initial photoreceptor specification at this developmental stage (Fig. 6E). Collectively, these findings suggest that *Tbx3* is required for the proper differentiation of RGCs and photoreceptors, which subsequently coordinate angiogenesis during early mouse postnatal retinal development.

### TBX3 is required for RGC differentiation

TBX5 is known to promote dorsoventral patterning (Peters and Cepko, 2002) and is involved in RGC differentiation (Sowden et al., 2001; Al-Khindi et al., 2022). As our RNAseq analysis showed *Tbx5* downregulation, we sought to validate this finding at the cellular level. In WT E13.5 retinas, TBX5 was robustly expressed by differentiating RGCs, spreading from the center towards the middle of the DCL. In cKO retinas, the number of TBX5^+^ RGCs was significantly reduced (Fig. 7A-C). To gain further insight into RGC differentiation, we examined the transient expression of Retinoid X Receptor γ (Hoover et al., 1998; Roberts et al., 2005) by RGC precursors in the DCL. Consistent with the β-tubulin and TBX5 data, cKO retinas exhibited a reduced number of RXRγ^+^ RGCs (Fig. 7A-B,D) when compared to WT littermates.

**Figure 7.**
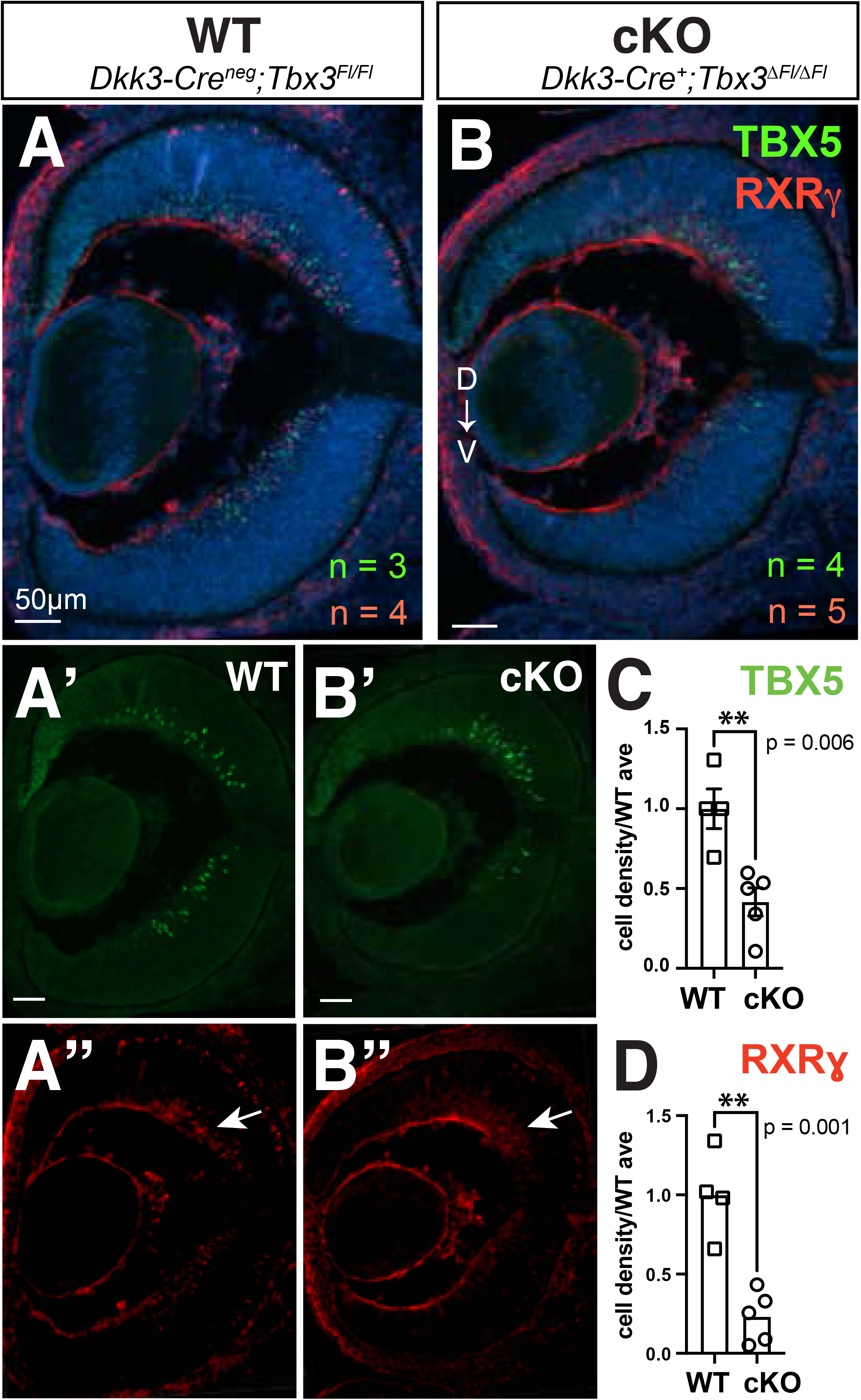
Early RGCs expressing TBX5 and RXRγ require TBX3. (A–B″) Retinal sections from WT (A–A″) and cKO (B–B″) E13.5 embryos stained for TBX5 (green) and RXRγ (red). points to emergent RGC layer in which RXRγ+ cells were counted. (C, D) Quantification of TBX5^+^ (C) and RXRγ^+^ (D) cells in the RGC layer. Data analyzed by Unpaired two-tailed Student’s t-test. Sample sizes (n) are color-coded green (TBX5) and red (RXRγ). Scale bar: 50 µm.

RGC fate commitment is canonically initiated by ATOH7, which subsequently induces the expression of *Isl1* and *Pou4f2* (Mu et al., 2005; Mu et al., 2008a; Gao et al., 2014; Wu et al., 2021). Since we observed reduced retinogenesis and the bulk RNA-seq analysis demonstrated a genetic signature of reduced ATOH7-ISL1-POU4F2-mediated RGC differentiation, we analyzed the expression of ATOH7, ISL1/2 and POU4F2 in E13.5 retinas.

Contrary to the hypothesis that reduced retinogenesis stems from diminished competence, *Tbx3* loss increased the number of ATOH7^+^ RPCs by ∼50% (WT: 1.0 ± 0.03; cKO: 1.57 ± 0.22; p = 0.05; Fig. 8A-C). Despite this increase in ATOH7, the density of ISL1/2^+^ RGC precursors was reduced by ∼15% in the cKO retinas (WT: 1.0±0.02; cKO: 0.87 ± 0.02; p=0.02; Fig. 8D-F). Similarly, POU4F2^+^ RGC precursors were reduced in cKO E13.5 retinas (WT: 0.98 ± 0.01 cKO: 0.81 ± 0.05; p = 0.02; Fig. 8G-I).

**Figure 8.**
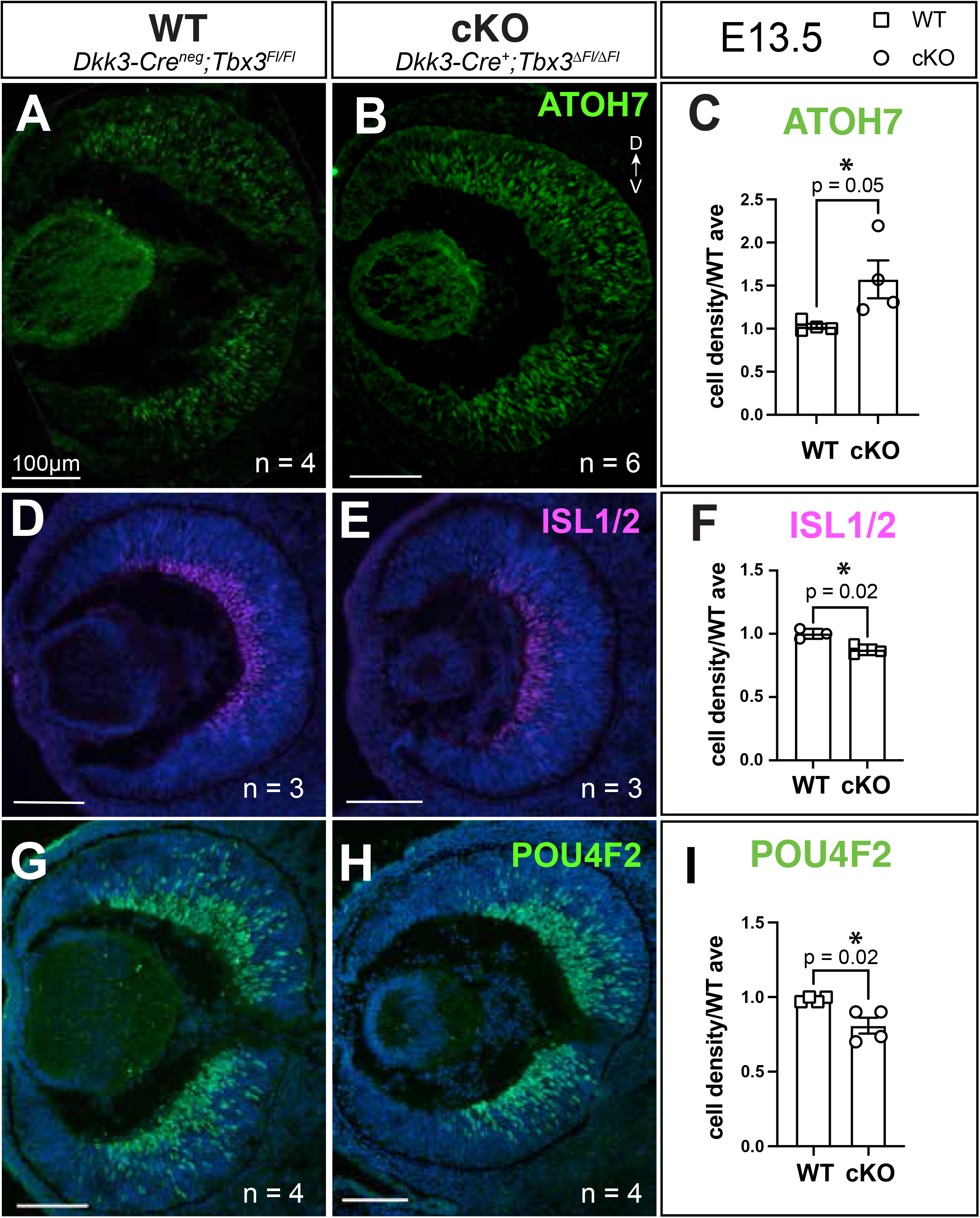
*Tbx3* loss increases ATOH7^+^ RPCs while reducing the known ATOH7 direct target genes, ISL1/2 and POU4F2, at E13.5. (A–C) Retinal sections from WT (A) and cKO (B) mice stained for ATOH7 (green) with quantification (C). (D–F) Retinal sections from WT (D) and cKO (E) mice stained for ISL1/2 (magenta) with quantification (F). (G–I) Retinal sections from WT (G) and cKO (H) mice stained for POU4F2 with quantification (I). Data analyzed by Unpaired two-tailed Student’s t-test. Scale bar: 100 µm.

The current model of RGC differentiation posits that perturbations in the ATOH7-ISL1-POU4F2 RGC differentiation program would cause a reduction in downstream targets, *Shh* and *Pou4f1* (Lyu et al., 2021), as observed in our transcriptomic analysis. To provide further evidence of the impact of *Tbx3* loss on the RGC differentiation program and further validate our RNA-Seq findings, we evaluated SHH and BRN3A expression. We observed a reduction in SHH (WT: 1.0 ± 0.06; cKO: 0.42 ± 0.10; p = 0.008) and BRN3A (WT: 1.0 ± 0.17; cKO: 0.40 ± 0.05; p = 0.03; Fig. 9A-D) cells in cKO E13.5 retinas, a phenomenon that was also present in the dorsal retina at P0 (SHH Dorsal WT: 1.0 ±0.03; cKO: 0.66 ± 0.10; p = 0.006; Ventral WT: 1.0 ± 0.05; cKO: 0.96 ± 0.08; p = 0.63; BRN3A Dorsal WT 1.0 ± 0.06; cKO 0.60 ± 0.09; p = 0.006; Ventral WT: 1.0 ± 0.08; cKO 0.83 ± 0.20; p = 0.42; Fig. 9E-H).

**Figure 9.**
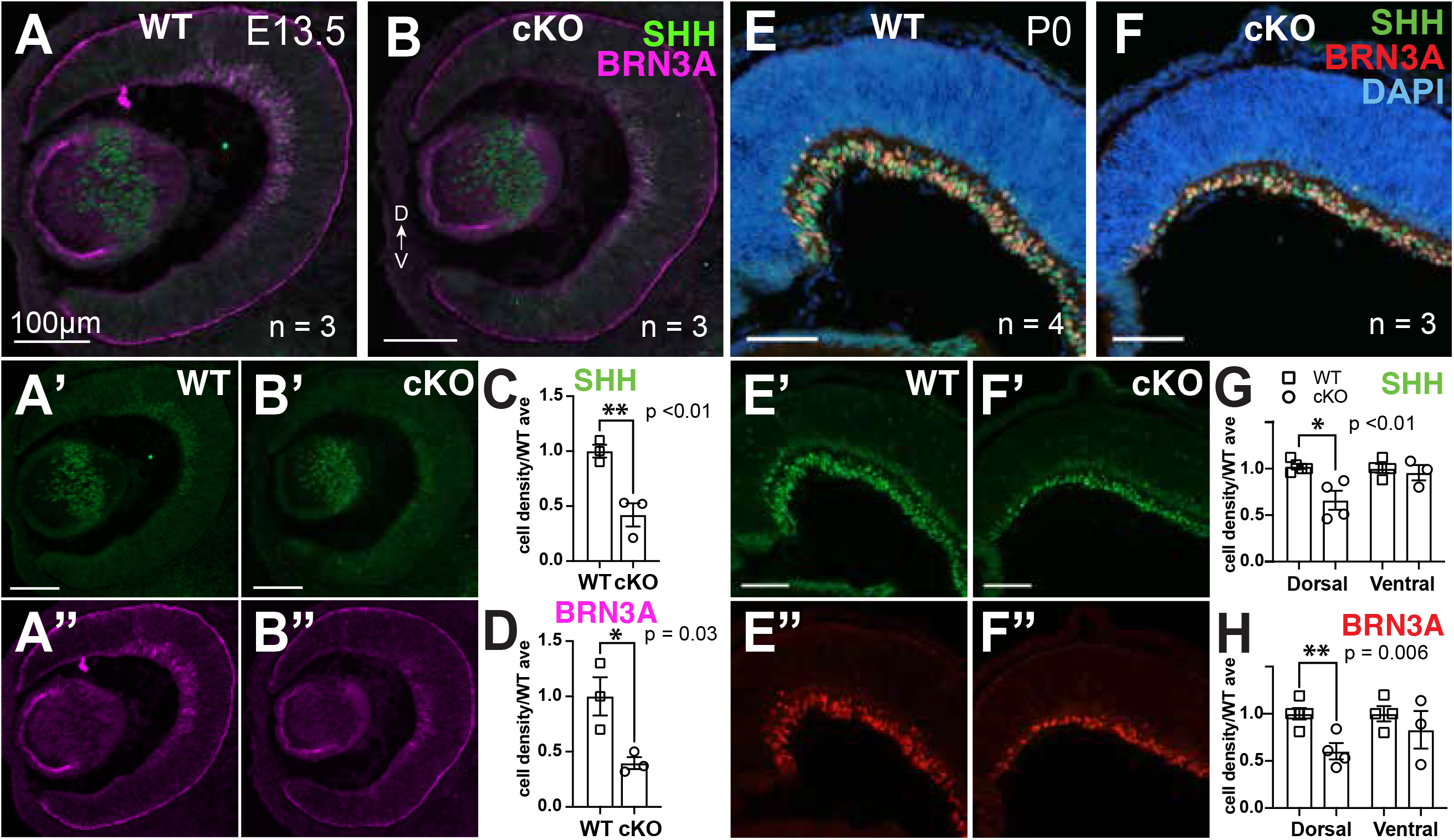
Canonical RGC-promoting factors require TBX3 in the embryonic retina, yet postnatally, they are only reduced in the dorsal eye. (A–D) E13.5 WT (A) and cKO (B) retinal sections co-stained for SHH (green) and BRN3A (magenta) with SHH^+^ (C) and BRN3A^+^ (D) cell quantifications. (E–H) P0 WT (E) and cKO (F) retinal sections co-stained for SHH and BRN3A with quantification showing dorsal-specific loss of SHH^+^ (G) and BRN3A^+^ (H) cells. Data analyzed by Unpaired two-tailed Student’s t-test. Scale bar: 100 µm.

Together, these data demonstrate that *Tbx3* loss reduces RGC differentiation by reducing the levels of ISL1/2^+^ and POU4F2^+^ RGC precursors at E13.5, leading to the consequential reduction of POU4F2^+^ RGCs and SHH in the postnatal retina. This indicates that TBX3 regulates the critical transition step from commitment to differentiation, thus explaining the observed retinogenesis defect.

### TBX3 is required for the formation of S-cones

*Tbx3* loss disrupts RGC differentiation, which subsequently stunts the formation of the superficial vascular plexus. Our previous study showed that the deep plexus was also defective upon *Tbx3* loss (Derbyshire et al., 2023). Deep vascular plexus forms in response to glutamatergic signals from photoreceptors (Biswas et al., 2024). Since our bulk RNA-Seq analysis showed reduced expression of key genes involved in cone development, we investigated whether *Tbx3* loss similarly affects cone development, potentially explaining the concurrent defects observed in the deep vascular plexus.

We first examined the formation of early photoreceptor precursors by staining E13.5 retinas for OTX2 and LHX4 (Nishida et al., 2003; Koike et al., 2007; Buenaventura et al., 2019). In WT retinas, OTX2 and LHX4 showed robust colocalization within the prospective outer nuclear layer (Fig. 10A). While this staining pattern was preserved in the cKO, the number of photoreceptor precursors was significantly reduced (Fig. 10A-C; WT: 1.0 ±0.07; cKO: 0.28 ± 0.05; p = ≤0.0001). As with RGC precursors, forcing retinal progenitor cells out of the cell cycle with DAPT *ex vivo* did not recover photoreceptor precursor formation in cKO retinas (Fig. S5B”,C”,E; WT: 1.0 ± 0.04; cKO: 0.09 ± 0.04, p=0.02).

**Figure 10.**
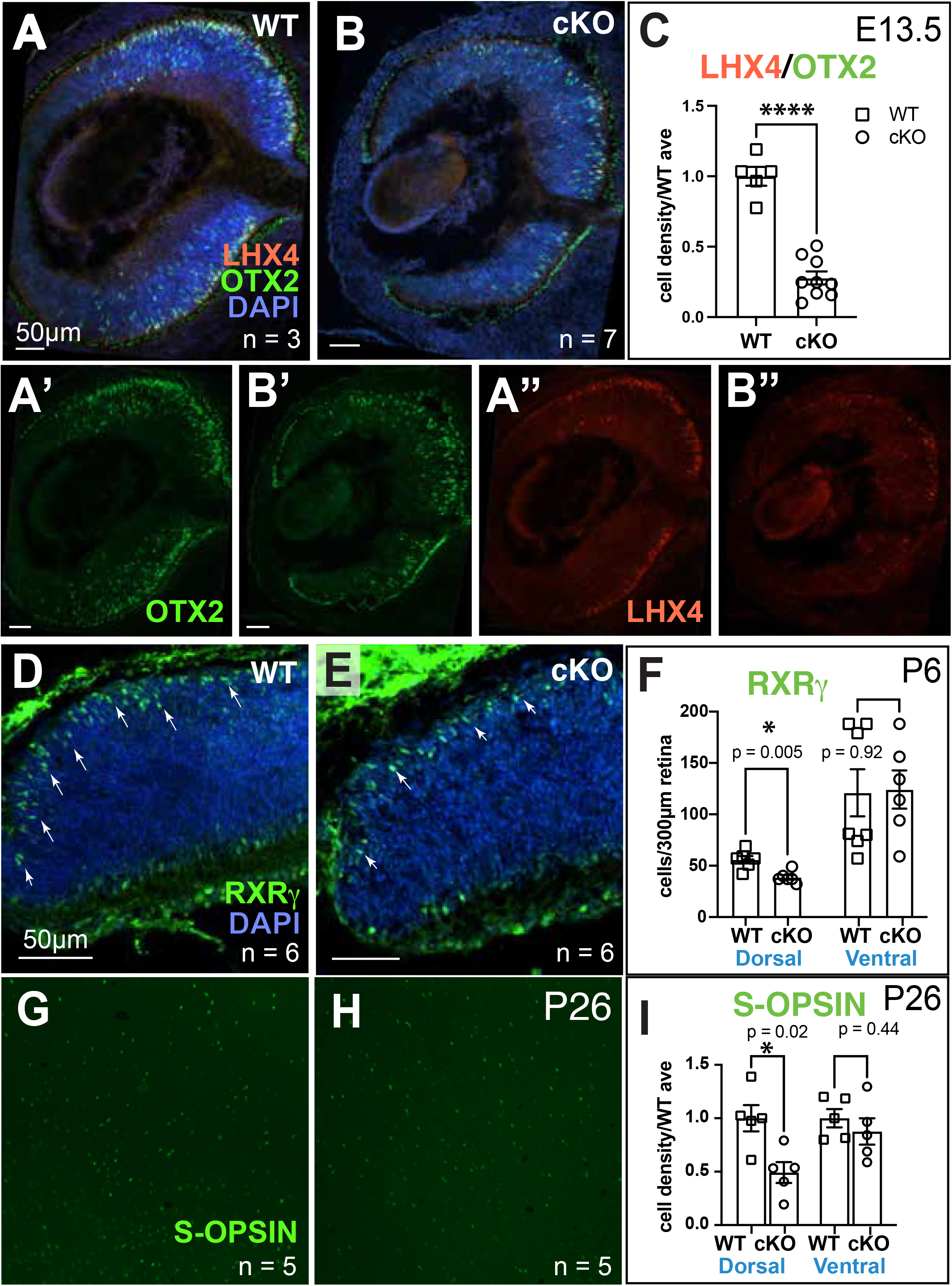
*Tbx3* is required for the formation of dorsal S-opsin^+^ cones. (A–C) E13.5 WT (A) and cKO (B) retinal sections stained for OTX2 (green; A’,B’) and LHX4 (red; A”,B”) with quantification of co-labeled cone precursors (C). (D–F) P6 WT (D) and cKO (E) dorsal retinal sections stained for RXRγ (green; arrows) and DAPI with quantification of the first 300 µm edge of the dorsal and ventral regions (F). (G–I) P26 WT (G) and cKO (H) close-up of dorsal retinal flat mounts stained for S-opsin and quantification of S-opsin+ cones at P26 (dorsal and ventral; I). Data analyzed by unpaired two-tailed Student’s t-test. Scale bars: as indicated.

To determine if this embryonic defect persisted postnatally, we focused on S-cone photoreceptors, the earliest cone subtype to differentiate (Szel et al., 1993). At P6, we stained retinas for RXRγ as a marker for developing S-cones (Roberts et al., 2005). By analyzing the cells in the outer nuclear layer, we observed a pronounced reduction in RXRγ^+^ cone numbers in the dorsal cKO retina, but the ventral cone population remained comparable between the genotypes (Fig. 10D-F; dorsal WT: 56 ± 3.6; cKO: 39 ± 2.3 cells/300µm retina, p = 0.005; ventral WT: 121 ± 23; cKO: 124 ± 19, p= 0.92). Consistent with these findings, analysis of P9 retinal flatmounts stained for S-opsin confirmed a significant overall reduction in S-cones in the cKO (Fig. S7A; WT: 1.0 ± 0.04; cKO: 0.57 ± 0.08, p= 0.02). Stratified analysis clarified that the primary effect of *Tbx3* loss was restricted to the dorsal retina (dorsal WT: 1.0 ± 0.04; cKO: 0.54 ± 0.06; p=0.005; Ventral WT: 1.0 ± 0.09; cKO: 0.88 ± 0.12; p=0.08). We observed by cross-sectional analysis that this defect was not due to a mislocalization of S-cones (data not shown), as observed in some models of ipRGC loss (Tufford et al., 2018). The S-cone loss was not transient: a 50% reduction in dorsal S-cones was maintained in mature P26 retinas (Fig. 10G-I; Fig. S7B,C; dorsal WT: 1.0 ± 0.12; cKO: 0.49 ± 0.10; p=0.012; Ventral WT: 1.0 ± 0.09; cKO: 0.88 ± 0.12; p=0.43). In contrast, when we examined M-cone density in P26 retinas using M-opsin staining, we found no difference between cKO and WT tissue (Fig. S7D-F; dorsal WT: 1.0 ± 0.22; cKO: 1.01 ± 0.24, p=0.97; ventral WT 1.0 ± 0.14 cKO 1.09 ± 0.07, p=0.61). This demonstrated a specific requirement for TBX3 in S-cone formation, but not M-cones, in the mature retina. Together, this data suggests that TBX3 is regulating the differentiation of a subset of RPCs required for both a subset of dorsal RGCs and S-cone formation.

## DISCUSSION

This work explored, for the first time, the role of TBX3 in the formation of RGCs and cone photoreceptors in the mouse retina and its potential consequence to retinal angiogenesis. We postulate that TBX3 acts as a fate determinant in early RPCs, based on the following evidence: (1) *Tbx3* expression as early as E9.0 in the mouse retina; (2) *Tbx3* loss leads to a defective transcriptomic signature of RGC and photoreceptor differentiation, which translates into reduced RGCs and cones in the embryonic and postnatal retina; (3) this TBX3-dependent effect likely takes place within the first RPCs to differentiate, as E12.5-E13.5 RGC birthdating showed reduced RGC precursor formation; (4) in the absence of *Tbx3*, there is a 40% increase in RPC proliferation (5) that when artificially inhibited *ex vivo*, does not recover the phenotype in cKO retinas. These early, essential effects of TBX3 in retinogenesis then impair angiogenesis, as loss of *Tbx3* within RPCs and not in mature RGCs or astrocytes impairs dorsal astrocyte migration and angiogenesis, specifically the superficial and deep vascular plexi, whose growth is known to be regulated by RGCs and photoreceptors, respectively.

### Role of TBX3 in retinogenesis

The spatiotemporal expression of *Tbx3* across diverse species suggests a conserved role for this transcription factor in specifying neural identity, particularly within the vision system. From an evolutionary perspective, *Tbx3* is expressed in some of the most rudimentary ocular structures, where a clear parallel exists between primitive eye types and the emergence of specific cell lineages (Kamijyo et al., 2015). In *Xenopus*, our group positioned Tbx3 at the forefront of the eye-field transcription factor (EFTF) cascade, acting as a downstream effector of the BMP inhibitor Noggin for retinogenesis (Zuber et al., 2003; Viczian et al., 2009; Motahari et al., 2016). In this study, we demonstrate that this requirement for TBX3-dependent retinogenesis persists in the mammalian retina.

The chronological development of retinal neuronal subtypes is driven by an intrinsic molecular clock within RPCs, yet how these broad temporal competence windows are translated into the precise specification of early-born lineages such as RGCs and cones remains an open question. Early and late RPC states are established by temporal identity factors (Blackshaw and Cayouette, 2025), which remain largely conserved in the *Tbx3* cKO retina, suggesting that TBX3 functions downstream of the primary clock or through a parallel mechanism to mediate early cell fates. Supporting this hierarchy, the early RPC factors MEIS1/2 directly bind the *Tbx3* promoter (Schindler et al., 2025) and their knockout shares similar phenotypes with our *Tbx3* cKO, with reduction in β-tubulin III, OTX2^+^ photoreceptor precursors, and RGCs, alongside a concomitant increase in apoptosis (Marcos et al., 2015). Mechanistically, the knockout of *Tbx3* or *Meis1/2* upregulate signaling molecules such as BMP7, BMPR1B, FZD1, and WNT7B, suggesting that a conserved MEIS-TBX3 module normally restrains these pathways to drive early retinogenesis. Furthermore, a non-cell-autonomous angiogenesis defect is also observed in *Meis1* mutants (Hisa et al., 2004), indicating that this module coordinates the broader developmental niche by integrating neurogenic and angiogenic signals necessary for retinal maturation.

Within this MEIS1/2-dependent early RPC framework, our findings indicate a direct role for TBX3 in the specification of RGCs and cones. We observed reduced RGC density in mature *Tbx3* cKO retinas, likely driven by the inefficient formation of RGC precursors. Our transcriptomic analysis showed downregulation of the canonical ATOH7-ISL1-POU4F2 program, suggesting that TBX3 diversifies the RGC program downstream of, or in parallel to the canonical pathway. This is illustrated by the loss of ipRGCs and direction-sensitive ganglion cells (DSRGCs) in *Tbx3* cKO retinas. ISL1 and POU4F2 operate upstream of TBR2 (Mao et al., 2008; Mu et al., 2008b), the lineage marker required for ipRGC specification (Zhang et al., 2014). *Tbx3* loss dismantles the ISL1/TBR2/OPN5 regulatory hierarchy necessary for these ancient neurons to form (Mao et al., 2008; Sajgo et al., 2017), leading to their absence in adult retinas (Derbyshire et al., 2023). Similarly, *Tbx3* loss interferes with the differentiation of DSRGCs, as we observed TBX5 downregulation at both the transcript and protein levels, accompanied by the differential expression of *Cnpe5, Mef2c*, and *Nptx2*, genes regulated by TBX5 in DSGCs (Al-Khindi et al., 2022).

Despite downregulation of the ATOH7-ISL1-POU4F2 differentiation program, ATOH7^+^ retinal RPCs increased upon *Tbx3* loss, an outcome explainable by two mechanisms. First, *Tbx3* loss may disrupt a TBX3-POU3F1 regulatory loop. POU3F1 represses *Tbx3* to drive neuroectodermal fate in stem cells (Zhu et al., 2014; Russell et al., 2015) and suppresses *Atoh7* and *Zic2* for the establishment of contralateral RGC fates (Fries et al., 2023). Reduced POU3F1 expression upon *Tbx3* cKO would relieve this repression, elevating ATOH7^+^ RPCs. However, our analysis demonstrated comparable levels of POU3F1 in WT and cKO E13.5 retinas (Fig. S4D-F) and no significant change in our RNA-Seq analysis. Second, this increase in ATOH7 may stem from loss of the SHH feedback. Differentiating RGCs secrete SHH to act on PTCH1^+^ RPCs. Reducing SHH promotes RGC differentiation at the expense of late fates (Zhang and Yang, 2001; Krueger et al., 2026). Thus, impaired RGC differentiation in *Tbx3* cKO retinas reduces SHH feedback, driving a compensatory surge in ATOH7^+^ RPCs and positioning TBX3 at the center of the balance between progenitor maintenance and RGC specification.

Parallel to its role in the RGC lineage, TBX3 is equally indispensable for the transcription factor cascade driving photoreceptor specification. This cascade begins with the photoreceptor fate commitment by OTX2 (Nishida et al., 2003; Emerson and Cepko, 2011; Emerson et al., 2013), which directly activates CRX (Nishida et al., 2003), while cooperating with BLIMP1 to repress bipolar or amacrine fates (Brzezinski et al., 2010; Katoh et al., 2010b; Goodson et al., 2020). Concurrently, LHX4 acts alongside OTX2 to regulate early RPC progression, ensuring the initial specification and expansion of the cone lineage (Buenaventura et al., 2019). NEUROD4 might act as a downstream bHLH factor promoting generic photoreceptor progression (Akagi et al., 2004), followed by the activation of nuclear receptors RXRγ and RORα to drive S-cone differentiation (Roberts et al., 2005; Fujieda et al., 2009). In *Tbx3* cKO retinas, we observe a distinct reduction in OTX2 and LHX4 progenitors, establishing that the cone progenitor pool is deficiently formed. This initial defect compromises the downstream signaling required to drive S-cone identity, as substantiated by the downregulation of *Crx, Blimp1*, and *NeuroD4* and the associated loss of RXRγ^+^ cones and mature S-cones in cKO retinas.

TBX3 has been reported as both, a repressor and an activator of gene expression. In analyzing published ChIP-seq data from other labs we observed that TBX3 is commonly found at the regulatory regions of the genes coding for the RGC factors EBF1, IRX5, IRX6, and POU4F1 and of the photoreceptor factors NEUROD4, BLIMP1, and RORα, which were differentially regulated in our transcriptomic analysis (Han et al., 2010; Zimmerli et al., 2020; Soussi et al., 2024). We postulate that by operating downstream of MEIS1/2 to unlock both the ATOH7-ISL1-POU4F2 RGC and the OTX/LHX4 cone progenitor cascades, TBX3 bridges broad temporal progenitor competence and the concrete execution of the early born retina neuron programs. This is substantiated by evidence showing that TBX3 overexpression at P0 induces amacrine and bipolar cell fates at the expense of rods, indicating that TBX3 induced an early cell fate, even in a developmental window poised to late cell fates (Lyu et al., 2021). Based on our observations, the absence of TBX3 leads to failed commitment of a subset of early RPCs, which persist in the cell cycle and ultimately undergo apoptosis. Alternatively, it is also possible that *Tbx3* loss also affects the maturation of RGC precursors, which would then explain the increase in apoptosis at E13.5 and reduced RGC density in mature retinas.

### A TBX Code in the Developing Mammalian Retina

The concept of a T-box code (a combinatorial regulatory system where specific factors dictate cell fate, analogous to the Hox code) has been characterized by the interplay between T-BET and TBR2 during lymphocyte differentiation (Nagel and Meyer, 2021). Our data suggests that a similar TBX code operates during retinal development. In the vertebrate retina, the expression of *Tbx2, Tbx3, and Tbx5* is strictly partitioned to define the dorsal-ventral axis across species, including chick, mice, humans, and *Xenopus* (Sowden et al., 2001; Wong et al., 2002). This code is highly responsive to BMP signaling, and in the absence of either BMP signaling or *Tbx* expression, dorsal retinal identity fails to establish, leading to significant patterning defects (Behesti et al., 2006).

Our findings demonstrate that TBX3 functions as a central scaffold within this mammalian retinal code, exhibiting a biological synergy comparable to the TBX code in the hematopoietic lineages. Upon *Tbx3* loss, we observed reduced expression of *Tbx2* and *Tbx5*, along with reduction in TBX5*^+^* RGCs. In addition, we previously demonstrated that *Tbx3* is co-expressed with a subset of TBR2^+^ cells and that its deletion leads to a 40% loss of these neurons in the adult retina (Derbyshire et al., 2023). These findings suggest that the TBX code governs both spatial patterning and the neurogenic competence of RPCs. Without the TBX3-mediated scaffolding of this code, RPCs fail to progress toward RGC or photoreceptor fates, identifying TBX3 as a requisite for the transition from multipotent precursor to differentiated neuron.

This model aligns with recent evidence in the chick retina, where cross-regulation between *Tbx2, Tbx3*, and *Tbx5* delimits the high acuity area (HAA). In that system, *Tbx* knockdown alters the expression of *Fgf8, Cyp26c1*, and *Cyp26a1*, causing defective HAA formation through dysregulated retinoic acid signaling (Ayten et al., 2026). In line with these findings, we also observed reduced expression of *Fgf8, Cyp26c1* and *Cyp26a1* upon *Tbx3* loss. Given the importance of these molecules to the HAA, it is tempting to speculate that TBX3 might be required signaling mechanisms that pattern the human fovea. Together, these results support the existence of a conserved TBX code in the developing retina that integrates positional cues with transcriptional mechanisms to drive precise patterning and cell differentiation.

### A conserved role for TBX3 in neurogenesis

The conserved role of TBX3 as a regulator of neurogenesis extends beyond the retina to the developing hypophysis. Expressed in the midline of the ventral diencephalon and Rathke’s pouch (the precursors of the neurohypophysis and adenohypophysis, respectively), *Tbx3* levels peak at E9.5, a period preceding cell fate commitment (Pontecorvi et al., 2008). Loss of *Tbx3* impairs the formation of both the neurohypophysis and adenohypophysis (Trowe et al., 2013). This defect is characterized by reduced expression of *Six3* and *Pax6*, which are essential for dorsal cell fate. Consequently, dorsal adenohypophysis patterning is lost, evidenced by the absence of progenitors for the TBX19^+^ (corticotrope/melanotrope) and POU1F1^+^ (lactotrope/somatostrope/thyrotrope) lineages. Similarly, in the infundibulum, *Tbx3* deficiency leads to dysregulated *Shh* expression and a loss of TBX2^+^ cells, ultimately halting neurohypophysis formation (Trowe et al., 2013). Evidence also suggest that *Tbx3* is required for the development of the arcuate nucleus in the hypothalamus. There, it is found within Kisspeptin, somatostatin, and melanocortin system neurons (AgRP/NPY and POMC), where it dictates cellular differentiation and identity. Collectively, the neurogenic roles of TBX3 in both sensory (retinal) and regulatory (hypophyseal and hypothalamic) regions of the CNS (Eriksson and Mignot, 2009; Sanz et al., 2015; Quarta et al., 2019; Ma et al., 2021) suggest it provides a fundamental transcriptional scaffold for early neurogenic programs. This indicates that this T-box family member was likely co-opted early in metazoan evolution to define the structural boundaries and functional identities of the CNS.

### Limitations

While this study establishes a conserved role for *Tbx3* in mammalian retinogenesis, investigating its role in mammalian eye field specification remains technically unfeasible because available eye-specific Cre lines activate only after this stage. Additionally, because we analyzed phenotypes at E13.5 (following knockout at ∼E10.5), some defects (like the reduction in OTX2^+^ and POU4F2^+^ precursors without transcriptomic changes) may reflect downstream effects rather than direct TBX3 activity. Nevertheless, the E13.5 data strongly corroborate the documented postnatal defects, validating this timepoint for analysis. Second, although our bulk RNA-Seq depth (40M reads) might have missed low-abundance DEGs, it successfully identified a defective ATOH7-ISL1-POU4F2 program that correlates with the RPC and differentiation defects validated via immunostaining. While we also detected differential regulation of dorsal-ventral genes, these spatial patterns require future histological validation. Because TBX3 can behave as both, repressor and an activator of gene expression, distinguishing its direct downstream targets from indirect secondary effects represents a critical step toward understanding its function in retinal patterning and retinogenesis. Elucidating these target networks is an ongoing objective of our laboratory. Finally, our genetic models definitively demonstrate that *Tbx3* is required within RPCs, rather than in either astrocytes or postnatal RGCs, to drive angiogenesis. Characterizing the precise molecular bridge linking TBX3-dependent retinogenesis and angiogenesis remains a focus of our lab. Ultimately, by establishing TBX3 as a conserved, cell-autonomous regulator of mammalian eye development, this work provides a framework for ongoing investigations into early ocular patterning and neurovascular signaling.

## ACKNOWLEDGMENTS

The authors would also like to thank Julianna Yager, Savannah Rutt, Chloe Becker, April Wright, Melissa Gonzalez and Lily Gravante for their technical assistance, Dr. Anne M. Moon for providing the *Tbx3*-floxed and *Tbx3^GR/+^* mouse lines, Galina Bachay and Professor William Brunken for providing the *Six3-Cre* mouse line, and the Developmental Studies Hybridoma Bank (DSHB).

## DATA AND RESOURCE AVAILABILITY

The Bulk RNA-seq data was deposited into the GEO database (GSE337180). All relevant data and details of resources can be found within the article and its supplementary information.

