## Supplementary material for "TBX3 is required for the differentiation of a subset of retinal ganglion and cone photoreceptor cells": text is in file

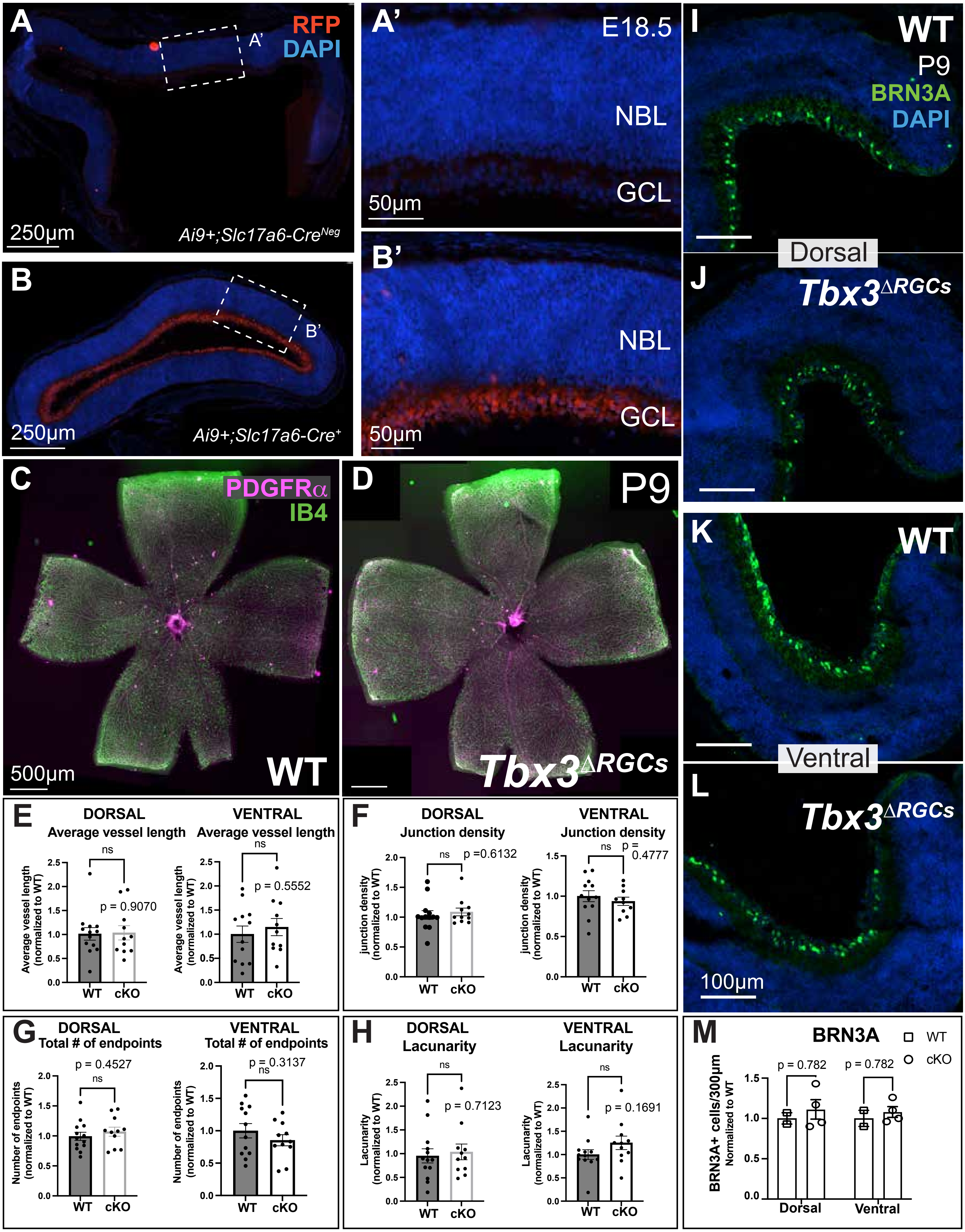


**Supplemental Figure S1.** ***Tbx3*** **loss in differentiated RGCs fails to affect retinal angiogenesis.**

(A-B) Representative images of E18.5 WT (A-A') and sibling *Ai9+;Slc17a6-Cre+* (B-B') retinal sections stained with anti-RFP antibody (red) and DAPI (nuclear stain) shows staining in the ganglion cell layer (GCL; B'), but not in the WT (A'). Scale bars as indicated in panels.

(C-D) Merged flat mount images of whole retinas from Fig. 2A,B. Scale bar, 500µm.

(E-F) Quantitation of vessel length (C), junction density (D), total number of endpoints (G) and lacunarity (H) when comparing the P9 *Tbx3*$^{\Delta RGC}$ to sibling WTs in the dorsal and ventral regions shown in Figure 2.

(I-L) Representative dorsal (I,J) and ventral (K,L) sections of P9 WT (I,K) and *Tbx3*$^{\Delta RGC}$ (J,L) retinas immunostained with anti-BRN3A antibody and DAPI. Scale bar, 100µm.

(M) Quantitation of the first 300µm of either dorsal or ventral BRN3A+ cells from central retina sections of WT (n=2); *Tbx3*$^{\Delta RGC}$ (n=4) where n is number of mice.

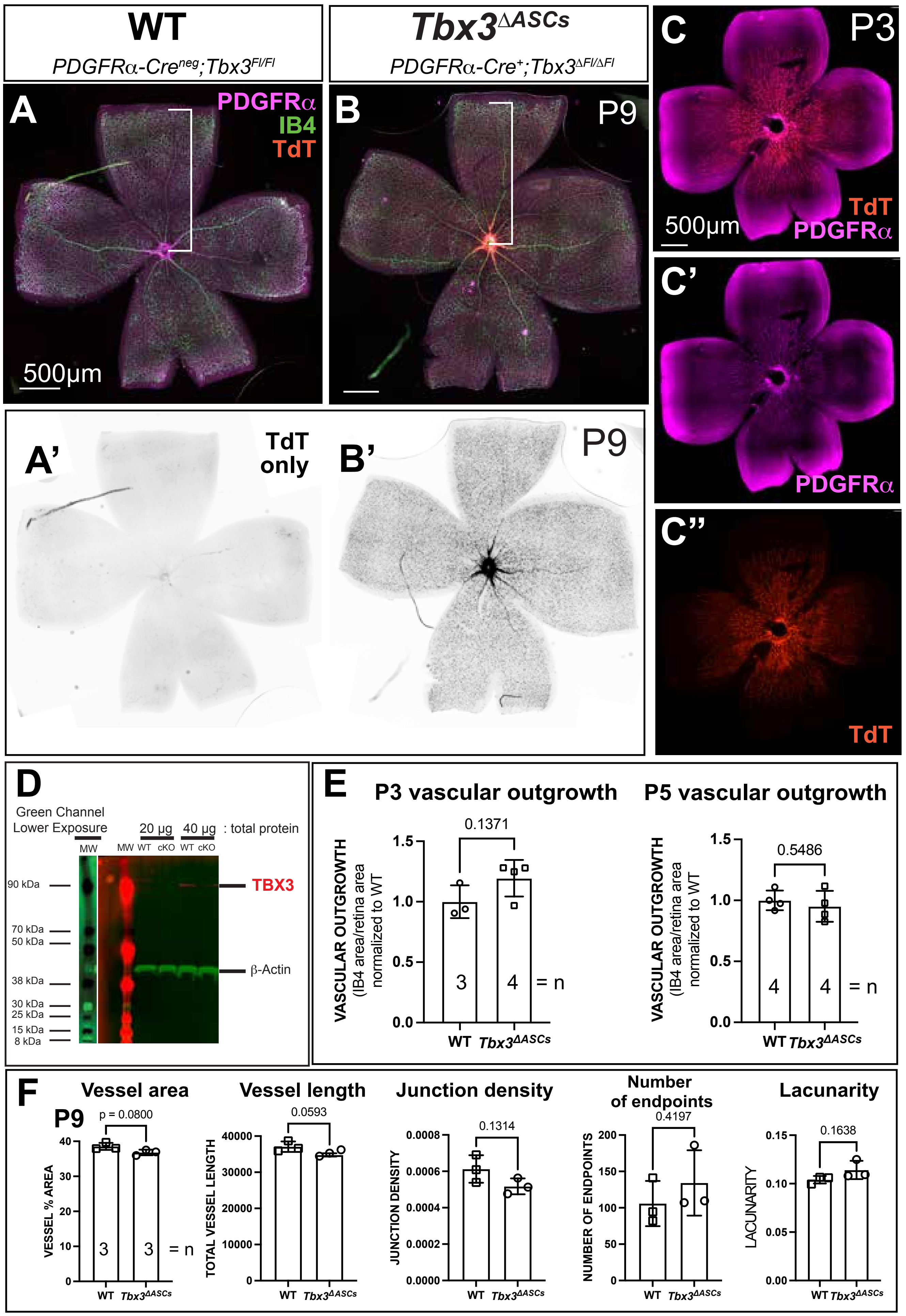


**Supplemental Figure S2. *Tbx3* loss in retinal astrocytes fails to affect retinal angiogenesis.**

(A-B) Retinal flat mounts at P9 from WT (A) and *Tbx3*$^{\Delta ASCs}$ (B) stained with anti-PDGFRα (astrocyte marker) and IB4 (blood vessel marker), while expression of Cre reporter is tdTomato fluorescent protein (TdT) that is visible without a stain. Inverted black and white image of TdT is visible in conditional knockout (B'), but not the WT (A') since the WT is Cre-negative.

(C) Retinal flat mount from a P3 *Tbx3*$^{\Delta ASCs}$ mouse (Cre+) stained with PDGFRα antibody (pink) and Cre-reporter TdT is visible (red, C). PDGFRα (C') and TdT (C") mark the same cells, suggesting successful Tamoxifen-induction of the Cre driver at P3, as well.

(D) TBX3 signal using western blot analysis is reduced in cKO (*Tbx3*$^{\Delta ASCs}$) sample compared to WT (compare 40µg protein samples cKO versus WT.

(E) Quantitation of vascular formation at P3 and P5 shows no significant change, although there is a trend toward a small increase at P3 which is not present by P5.

(F) Quantitation of P9 vessel area, vessel length, junction density, total number of endpoints, and lacunarity when comparing the *Tbx3*$^{\Delta ASCs}$ to sibling WTs in Figure 3.

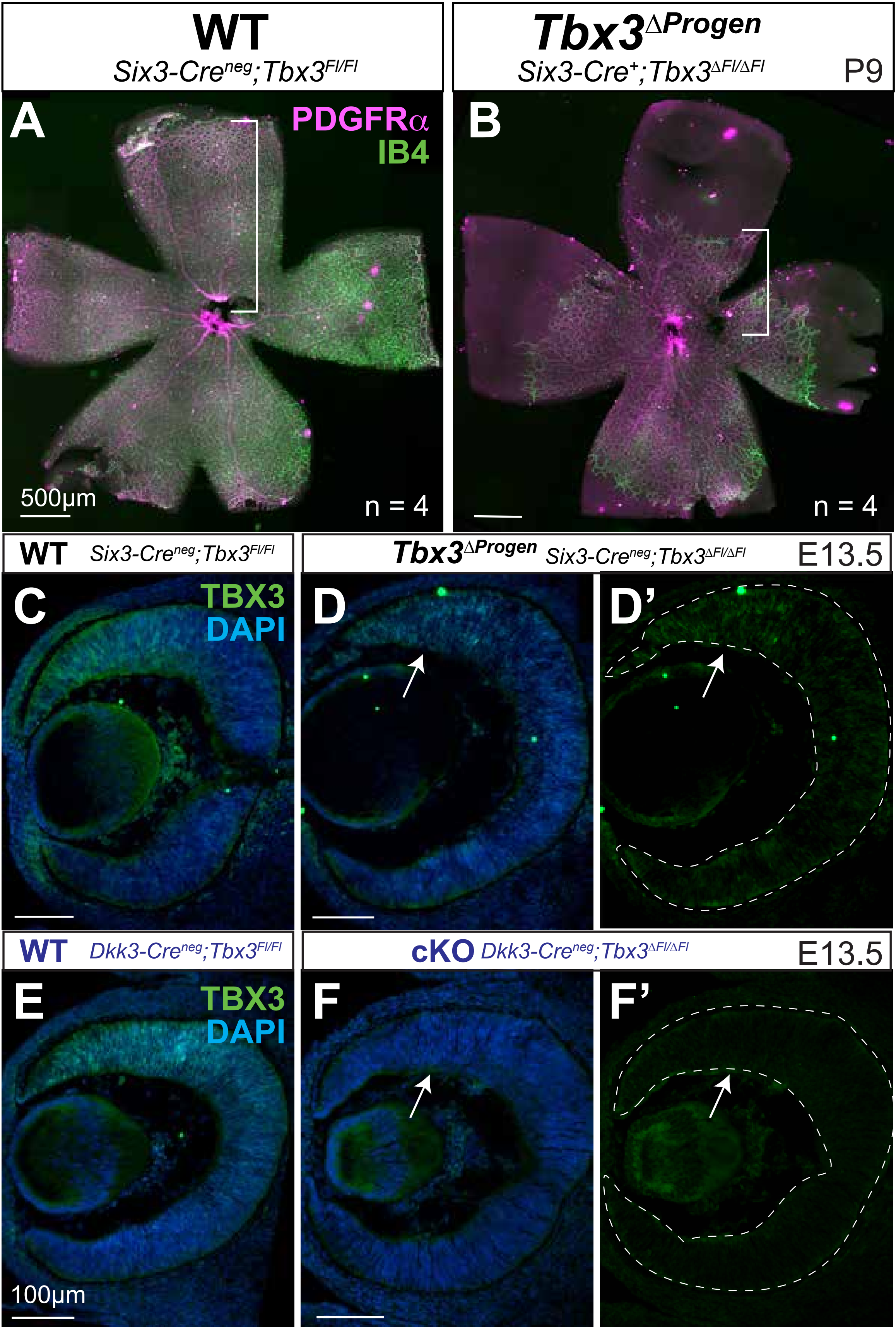


**Supplemental Figure S3. TBX3 removal with *Six3-Cre* causes a hypovascular phenotype, yet this Cre driver fails to fully remove TBX3.** (A-B) Merged retinal flat mounts from Fig. 4A,B immunostained with anti-PDGFRa (pink) and IB4-FITC (green). Brackets mark the retinal center to the edge of the retinal vascular growth. (C-F') Retinal E13.5 sections from two different conditional knockout lines immunostained with anti-TBX3 antibody (green) and DAPI (blue). (C) Stereotypical TBX3 dorsal expression in wildtype is only partially lost in (D, D') sibling *Tbx3*$^{\Delta Progen}$ retina (arrow), while, in a second conditional knockout of *Tbx3,* TBX3 expression in (E) wildtype mice is absent from the retina of (F,F') sibling *Dkk3-Cre;Tbx3*$^{\Delta Fl/\Delta Fl}$ mice (arrows). Note that TBX3 is present in vascular cells in F' as *Dkk3-Cre* targets retinal progenitors and astrocytes. Scale bars indicate 500μm in A,B and 100μm in C-F'. Dashed lines outline edge of the neural retina of cKO mice.

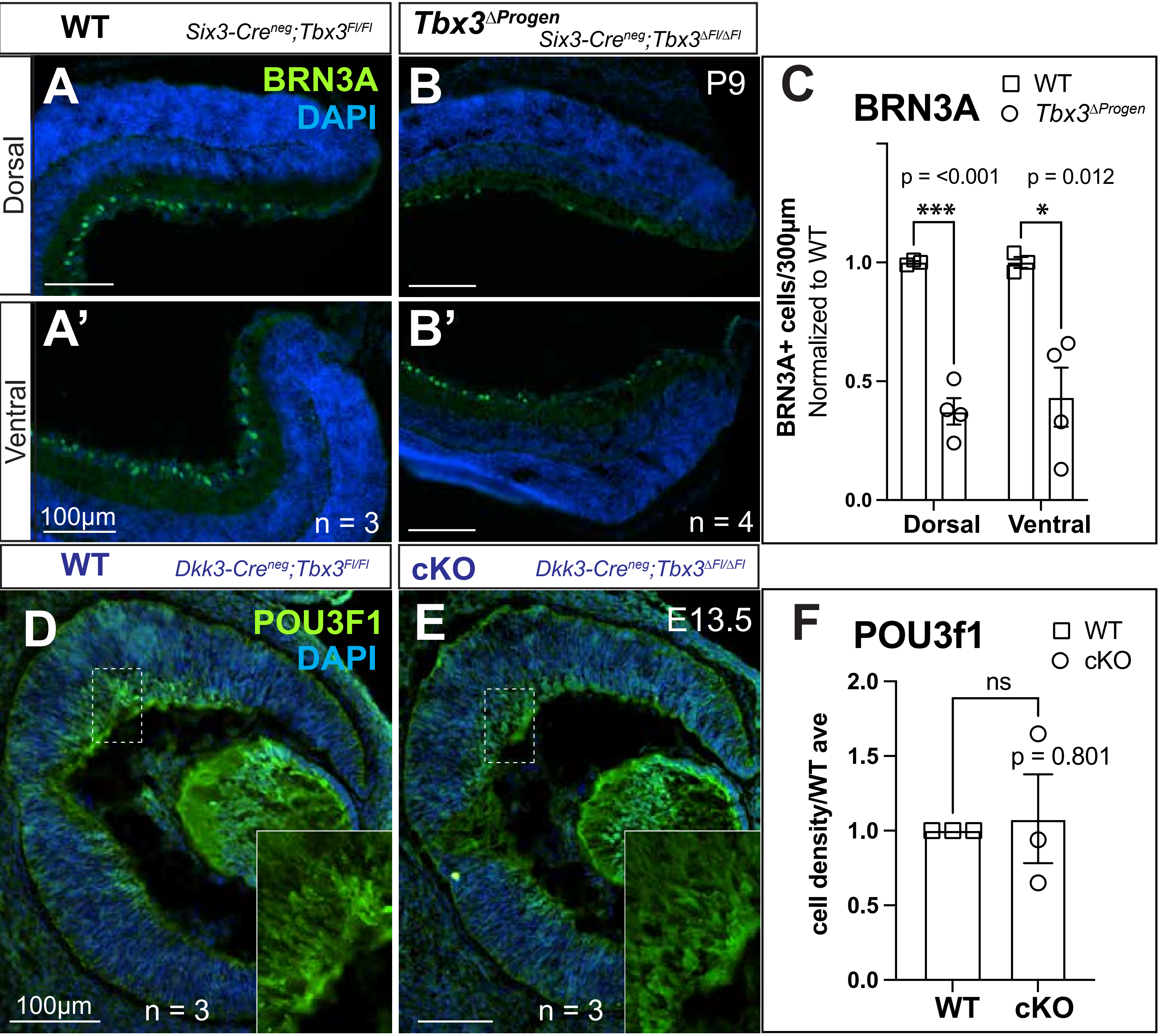


**Supplemental Figure S4. TBX3 is required for BRN3A expressing retinal ganglion cells, but not POU3F1 expressing cells.** (A-B') Representative sections from dorsal (A,B) and ventral (A',B') P9 sibling WT and *Tbx3*$^{\Delta Progen}$ retinas immunostained with anti-BRN3A antibody(green) and nuclear stain, DAPI (blue). Scale bars, 100μm; number of mice, n.
(C) Quantitation of BRN3A+ cells counted in both dorsal and ventral regions.
(D-E) Retinal E13.5 sections from WT (D) and sibling cKO retina (E) immunostained with anti-POU3F1 antibody (green) and DAPI (blue). Dashed lines indicates area in inlay, which is only POU3F1 staining. Scale bars, 100μm; number of mice, n.
(F) Quantitation of the POU3F1+ cells in WT and cKO retinas. Statistical comparison shows that they are not significantly (ns) changed.

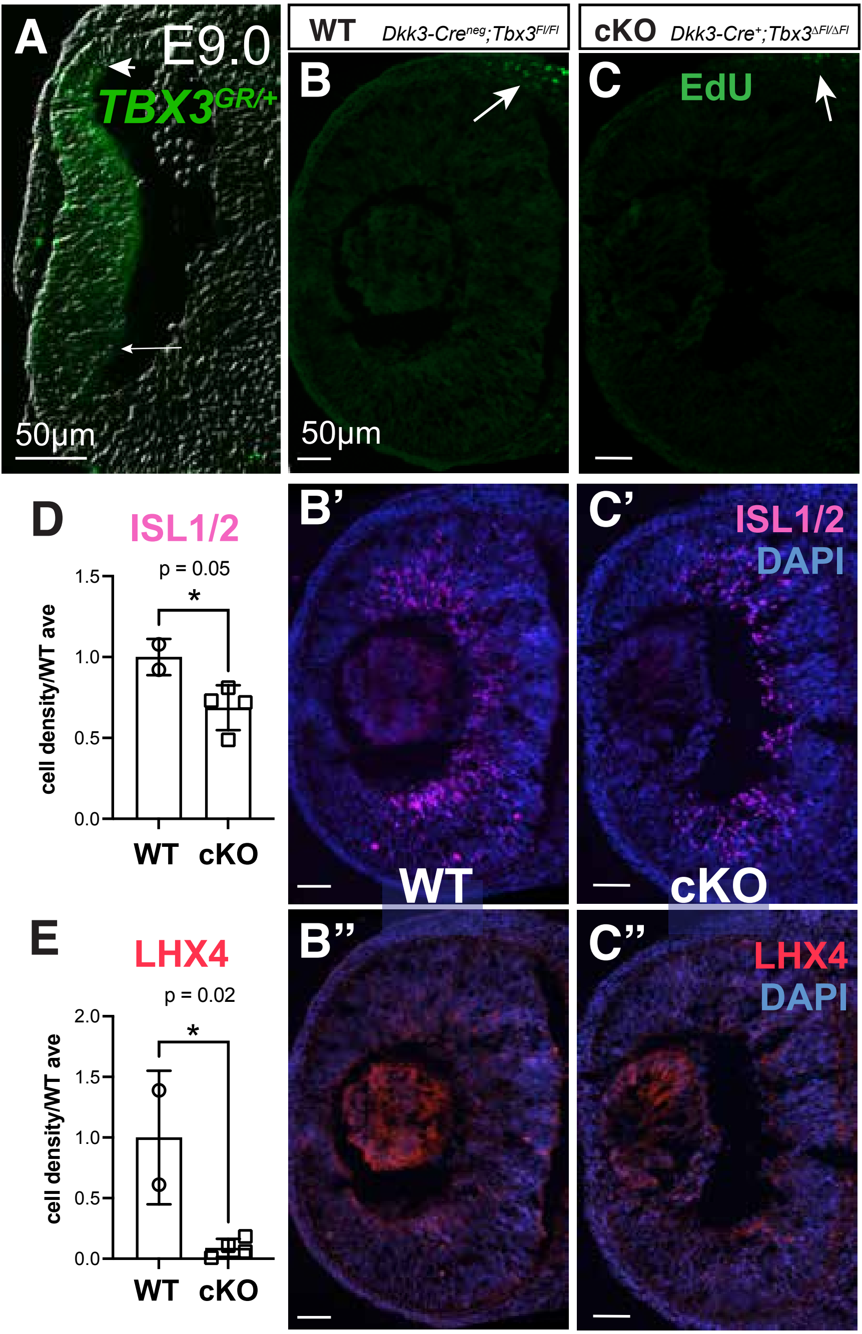


**Supplemental Figure S5. Forcing RPCs out of the cell cycle fails to recover RGC or cone precursor cell fate.** (A) Expression of Tbx3 occurs early, as indicated by GFP expression in E9.0 optic vesicle section of an embryo with GFP knocked-into the *Tbx3* gene locus (*Tbx3*$^{GR/+}$). Overlay of GFP onto brightfield image. Arrows point to edges of GFP staining.

(B-C") Representative retinal sections of WT (B-B") and cKO (C-C") cultured eye cups treated with Notch pathway inhibitor, DAPT, and EdU (green) for 24 h. EdU detection (B-C, green), anti-ISL1/2 (B'-C', pink) and anti-LHX4 antibodies (B"-C", red) and nuclear marker, DAPI (blue) stained sections. Arrows in B and C point to EdU$^{+}$ cells outside the retina. Scale bars are 50µm.

(D-E) Quantitation shows no recovery of the ISL1/2$^{+}$ cells when comparing cKO to WT (C). Likewise, cone precursors (LHX4$^{+}$ cells) remain significantlly reduced in the cKO retinas compared to WT (D).

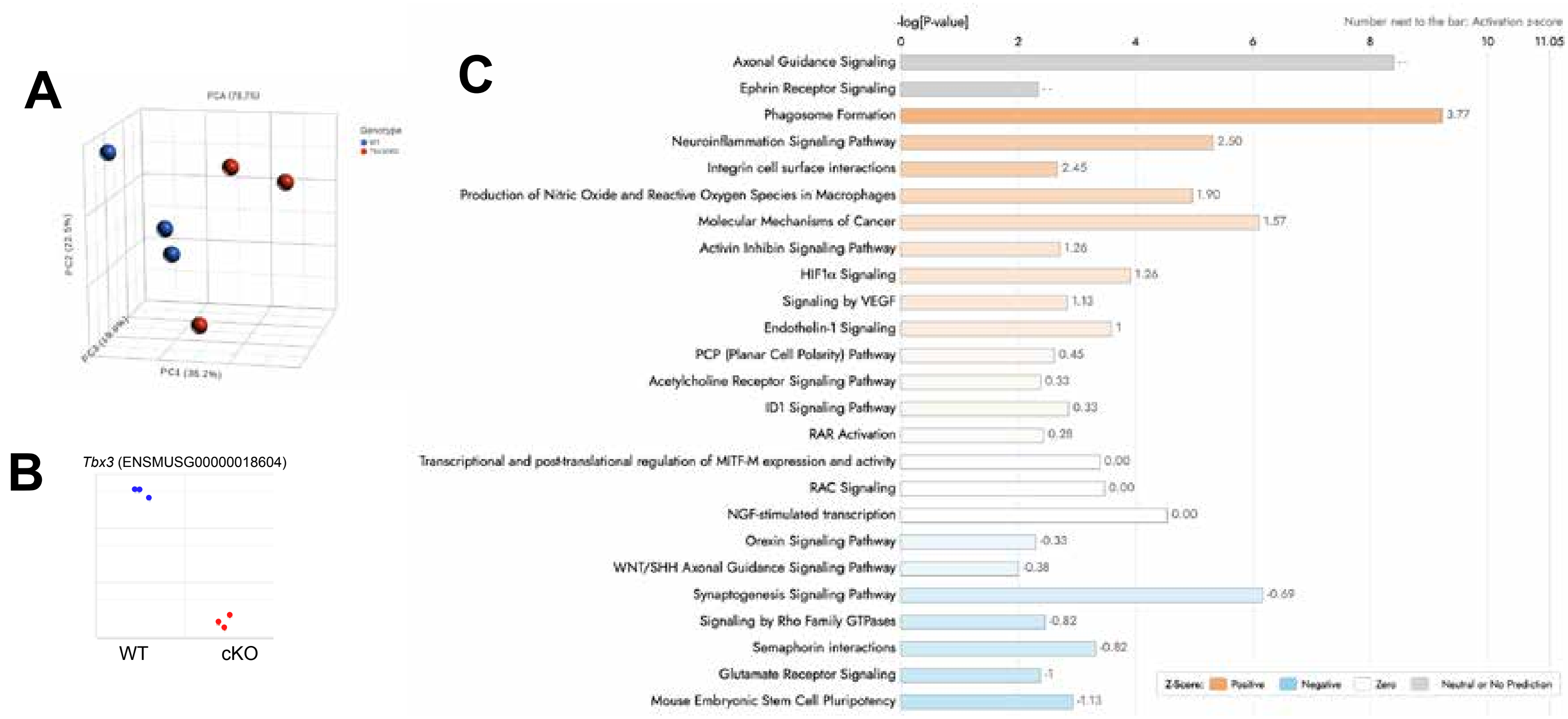


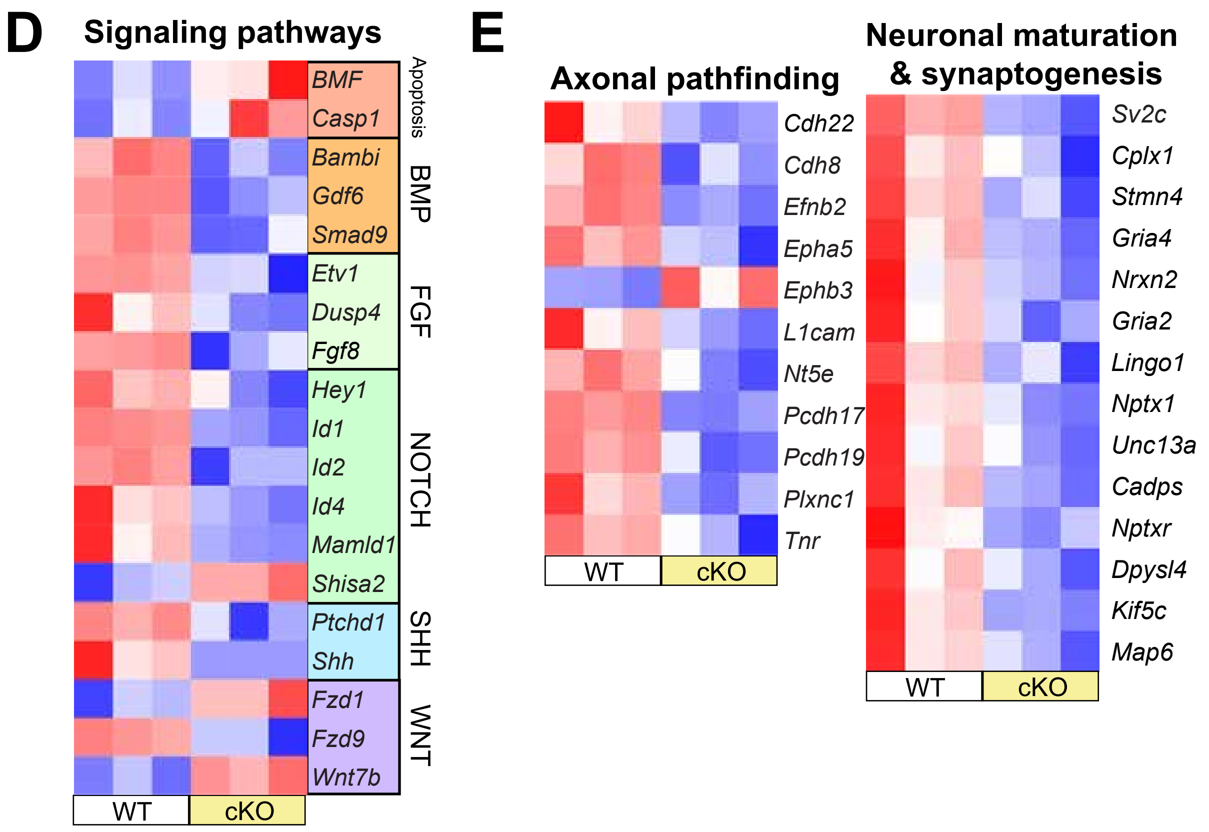


**Supplemental Figure S6. TBX3 is required for retinal neural development.**
(A) 3D PCA graph of E13.5 WT (blue) and cKO (red) retinal samples.
(B) Log plot of *Tbx3* mRNA samples shows a statistically significant fold-change between WT (blue) and cKO (red) samples.
(C) Canonical pathways identified by IPA analysis were predicted to be upregulated (orange) or downregulated (blue) or simply affected (gray) by *Tbx3* loss. Each is given an ‘activation score’ shown to the right of each bar. Pathways are listed on the left.
(D-E) Pathways we identified were significantly changed with a FDR score ≤ 0.05.

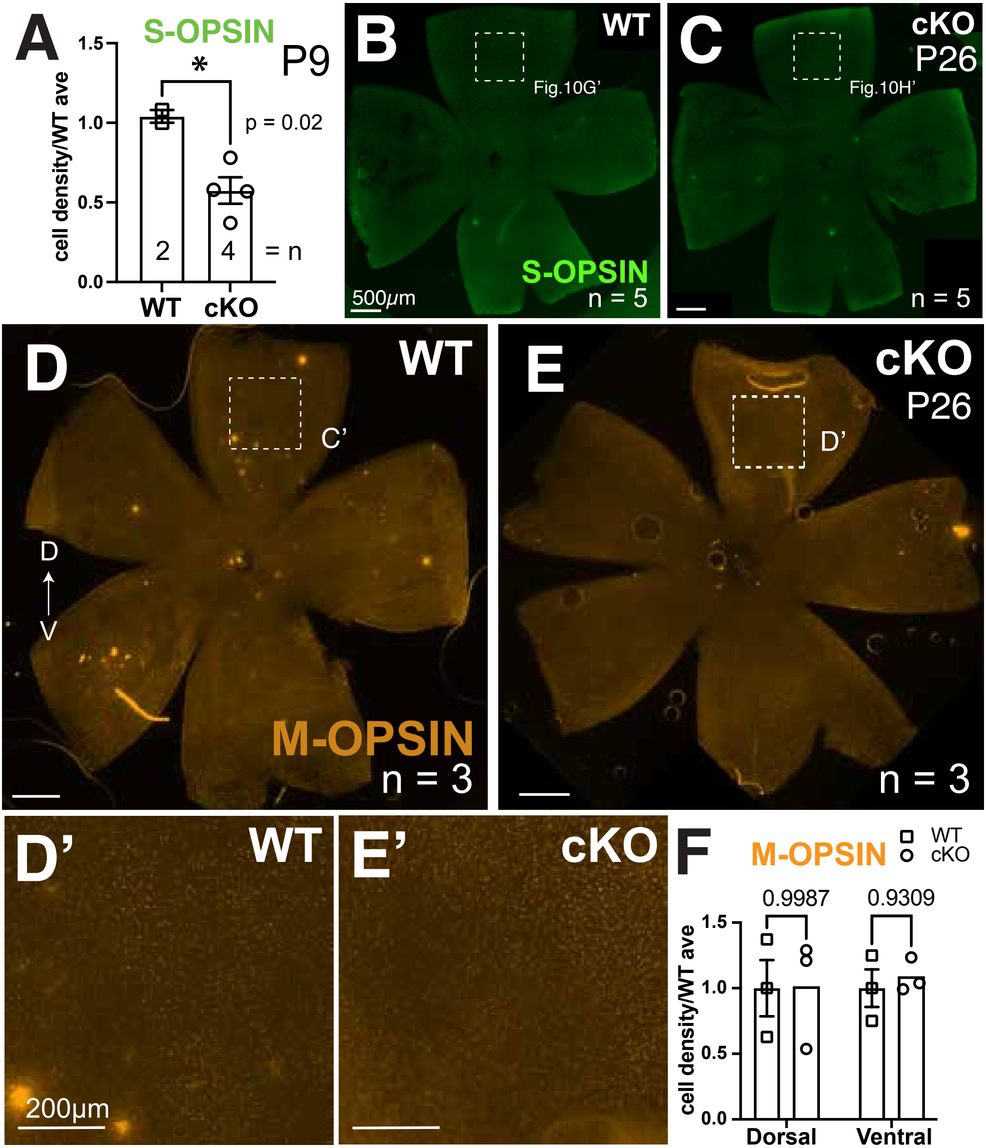


**Supplemental Figure S7. S-Cones, but not M-Cones, are not affected by *Tbx3* loss.**

(A) Quantitation of density of P9 S-Opsin+ cells normalized to the average WT density in retinal flat mounts.

(B-C) Retinal flat mounts of P26 WT and cKO (*Dkk3-Cre;Tbx3*$^{\Delta Fl/\Delta Fl}$) immunostained with anti-S-Opsin antibody (green).

(D-E') Immunostained P26 retinal flat mounts with anti-M-Opsin antibody (orange).

(E) Quantitation of density of M-Opsin+ cells counted normalized to the average WT density in dorsal and ventral areas. Number of mice is 'n'; scale bars are 500μm for A-D and 200μm for C'-D'.

Dashed boxes mark the close-up images in panels indicated. Dorsal (D) is up in all images.

**Supplemental Table S1. Antibodies and experimental conditions for immunostaining.**

| Target | Species raised | Company | Catalog # | Lot # | Blocking conditions | 1° Ab conc | 2° Ab co. & cat # | Antibody ID | PMID |
|---|---|---|---|---|---|---|---|---|---|
| **β-Tubulin** | Mouse | Covance | MMS-435P | | 1X PBS, 5% HIGS, 0.5% Triton | 1:400 | Invitrogen A21137 | N/A | 12798297 33428890 |
| **Atoh7** | Rabbit | Novus Biologicals | NBP1-88639 | A106752 | 1X PBS, 5% HIGS, 0.5% Triton | 1:300 | Invitrogen A21206 | N/A | 29977079 |
| **Brn3a** | Guinea Pig | Synaptic Systems | 411004 | | 1X PBS, 10% HIHS, 0.3% Triton | 1:1000 | Jackson Lab 706-585-148 | AB_2800538 | 37590135 |
| **Brn3b** | Goat | Santa Cruz | sc-6026 | D1012 | 1X PBS, 10% HIGS, 0.5% Triton | 1:250 | Invitrogen A34849 | AB_673441 | 21935940 27957530 |
| **Cleaved caspase-3** | Rabbit | Cell Signaling | 9664 | 21;ref 6/2018 | 1X PBS, 10% HIDS, 0.5% Triton | 1:400 | Invitrogen A31572 | AB_331439 | |
| **Islet 1/2** | Mouse | DSHB | 39.4D5 | N/A | 1X PBS, 10% HIDS, 0.3% Triton | 1:100 | Invitrogen A31570 | AB_2314683 | 1350865 31313880 |
| **Ki67** | Rat | eBioscience/ Fischer | 14-5698-82 | 2496198 | 1X PBS, 10% HIDS, 0.3% Triton | 1:3000 | Invitrogen A11077 | N/A | 33960384 |
| **Lhx4** | Rabbit | Proteintech | 11183-1 AP | 000409077 | 1X PBS, 5% HIHS, 0.5% Triton | 1:250 | Invitrogen A21206 | N/A | 35388544 |
| **Lim1/2** | Mouse | DSHB | 4F2 | N/A | 1X PBS, 10% HIDS, 0.5% Triton | 1:30 | Invitrogen A31570 | AB_531784 | 7528105 |
| **M-opsin** | Rabbit | Millipore | AB5405 | LV1442442 | 1X PBS, 5% HIGS, 0.3% Triton | 1:200 | Invitrogen A21206 | N/A | 2643825 |
| **Otx2** | Goat | R&D Systems | AF1979 | KNO041506A | 1X PBS, 5% HIHS, 0.5% Triton | 1:250 | Invitrogen A34849 | N/A | 29148976 |
| **Pax6** | Mouse | DSHB | Pax6 | N/A | 1X PBS, 10% HIGS, 0.4% Triton | 1:50 | Invitrogen A31570 | AB_528427 | 12648492 18626943 |
| **Pou3f1** | Rabbit | GeneTex | GTX134063 | 44539 | 1X PBS, 10% HIHS, 0.3% Triton | 1:250 | Invitrogen A31572 | AB_2887197 | 29791852 |
| **RXRγ** | Mouse | Santa Cruz | sc-365252 | D0623 | 1X PBST, 10% HIGS, 0.1% Triton | 1:25 | Invitrogen A31570 | N/A | 33712461 |
| **Shh** | Goat | R&D Systems | AF464 | | 1X PBS, 10% HIHS, 0.3% Triton | 1:500 | Invitrogen A11055 | AB_35537 | |
| **S-opsin** | Rabbit | Millipore | AB5407 | LV16557457 | 1X PBS, 5% HIGS, 0.3% Triton | 1:400 | Invitrogen A21206 | N/A | 25228773 |
| **Tbx5** | Rabbit | Novus Biologicals | NBP1-83237 | D115011 | 1X PBST, 10% HIGS, 0.1% Triton | 1:500 | Invitrogen A21206 | N/A | 33075808 |
| **Iba1** | Rabbit | Fujifilm | 019-19741 | LDQ3810 | 1X PBST, 10% HIHS, 0.3% Triton | 1:1000 | Invitrogen A21206 | AB_839504 | 8713135 |

**Supplemental Table S2. Ingenuity Pathway Analysis (IPA) of differentially regulated pathways when comparing WT and Tbx3 cKO E13.5 retinas**

| Activation z-score | | 3.77 | 2.5 | 2.45 | 1.9 | 1.57 | 1.26 | 1.26 | 1.13 | 1 |
|---|---|---|---|---|---|---|---|---|---|---|
| **Axon Guidance** | **Ephrin Receptor Signaling** | **Phagosome Formation** | **Neuroinflammation** | **Integrin cell surface interactions** | **Production of NO and ROS in Macrophages** | **Molecular Mech of Cancer** | **Activin Inhibin Signaling Pathway** | **HIF1α Signaling** | **Signaling by VEGF** | **Endothelin-1 Signaling** |
| ADAMTS15 | EFNB2 | ADGRB2 | CD86 | ITGA2B | IKBKE | ADGRB2 | EGLN3 | EGLN3 | NCF1 | HMOX1 |
| ADAMTS16 | EPHA5 | ADGRE1 | CSF1R | ITGAM | IRF8 | ADGRE1 | IGHM | HMOX1 | NCF4 | MAPK10 |
| ARPC1B | EPHB3 | ADRB2 | CX3CR1 | ITGB2 | Lyz1/Lyz2 | ADRB2 | IKBKE | HSPA1A | NCKAP1L | PIK3R5 |
| BMP7 | ITGA2B | ARPC1B | FZD1 | SPP1 | MAPK10 | BMP7 | MAPK10 | MET | PAK3 | Pla2g5 |
| EFNB2 | ITGAM | C3 | H2-T-ps/H2-T23 | VTN | NCF1 | BMPR1B | NGFR | NCF1 | PRKCB | PLCB2 |
| EPHA5 | ITGB2 | C3AR1 | HMOX1 | | NCF4 | C3AR1 | PAX2 | PIK3R5 | VAV1 | PLCL1 |
| EPHB3 | PAK3 | C5AR1 | IKBKE | | NGFR | C5AR1 | PIK3R5 | PRKCB | | PLD5 |
| FZD1 | RAC2 | C5AR2 | KCNJ3 | | PIK3R5 | C5AR2 | SMAD9 | RAC2 | | PRKCB |
| FZD9 | | CLEC7A | MAPK10 | | PRKCB | CNR1 | TLR7 | RORC | | PTGER2 |
| ITGA2B | | CNR1 | NCF1 | | RAC2 | CSF2RB | | SAT1 | | PTGS1 |
| ITGAM | | CX3CR1 | PIK3R5 | | S100A8 | CTNNA2 | | TF | | |
| ITGB2 | | F2RL1 | Pla2g5 | | SPI1 | CX3CR1 | | | | |
| KEL | | FCER1G | RAC2 | | | EDN1 | | | | |
| L1CAM | | Fcgr3 | SLC6A11 | | | F2RL1 | | | | |
| LINGO1 | | FZD1 | SYK | | | FZD1 | | | | |
| MET | | FZD9 | TLR7 | | | FZD9 | | | | |
| NGFR | | GRM5 | TREM2 | | | GRM5 | | | | |
| PAK3 | | GRM7 | | | | GRM7 | | | | |
| PAPPA2 | | HMOX1 | | | | HEY1 | | | | |
| PIK3R5 | | IGHM | | | | ID1 | | | | |
| PLCB2 | | ITGA2B | | | | ITGA2B | | | | |
| PLCL1 | | ITGAM | | | | ITGAM | | | | |
| PLXNC1 | | ITGB2 | | | | ITGB2 | | | | |
| PRKCB | | PAK3 | | | | MAPK10 | | | | |
| RAC2 | | PIK3R5 | | | | PAK3 | | | | |
| SHH | | Pla2g5 | | | | PIK3R5 | | | | |
| Tuba4a | | PLD5 | | | | PLCB2 | | | | |
| WNT7B | | PRKCB | | | | PRKCB | | | | |
| | | PTGER2 | | | | PTGER2 | | | | |
| | | RAC2 | | | | RAC2 | | | | |
| | | RASGRP1 | | | | RASGRP1 | | | | |
| | | SYK | | | | SHH | | | | |
| | | TLR7 | | | | SMAD9 | | | | |
| | | VAV1 | | | | | | | | |

| 0.45 | 0.33 | 0.33 | 0.28 | 0 | 0 | 0 | -0.33 | -0.38 | -0.69 | -0.82 | -0.82 | -1 | -1.13 |
|---|---|---|---|---|---|---|---|---|---|---|---|---|---|
| **PCP (Planar Cell Polarity) Pathway** | **Acetylcholine Receptor Signaling Pathway** | **ID1 Signaling pathway** | **RAR Activation** | **Transcriptional and post-translational regulation of MITF-M expression and activity** | **RAC Signaling** | **NGF-stimulated Xcription** | **Orexin Signaling Pathway** | **WNT/SHH Axonal Guidance Signaling Pathway** | **Synapto-genesis Signaling Pathway** | **Signaling by Rho Family GTPases** | **Semaphorin interactions** | **Glutamate Receptor Signaling** | **Mouse embryonic stem cell pluripotency** |
| FZD9 | CHRNB1 | BMPR1B | Aldh1a1 | ID1 | CYFIP2 | EGR3 | CACNG5 | FZD1 | ARPC1B | CDH22 | MET | GRIA4 | FZD9 |
| MAPK10 | HMOX1 | CHRNB1 | ALDH1A3 | POU3F2 | ITGA2B | ID1 | CNR1 | FZD9 | CDH22 | CDH8 | PAK3 | GRID1 | ID1 |
| ROR2 | NGFR | EGR1 | CSF2RB | TBX3 | ITGAM | ID2 | KCNJ3 | MAPK10 | CDH8 | ITGA2B | PLXNC1 | GRM5 | ID2 |
| WNT7B | PIK3R5 | ID1 | CYP26A1 | TFEB | ITGB2 | ID4 | PIK3R5 | PIK3R5 | CPLX1 | ITGAM | PTPRC | GRM7 | ID4 |
| | PLCB2 | NGFR | FGF8 | TFEC | NCF1 | LYL1 | Pla2g5 | SHH | EFNB2 | ITGB2 | TREM2 | | PIK3R5 |
| | PLCL1 | PIK3R5 | IGFBP3 | | PAK3 | | PLCB2 | SHISA2 | EPHA5 | MAPK10 | | | SMAD9 |
| | PRKCB | S100A9 | MAPK10 | | PIK3R5 | | PLCL1 | WNT7B | EPHB3 | NCF1 | | | |
| | TH | SHH | MPPED2 | | RAC2 | | PRKCB | | GRIA2 | PAK3 | | | |
| | | SMAD9 | NR2F1 | | | | SMAD9 | | GRIA4 | PIK3R5 | | | |
| | | | PIK3R5 | | | | | | GRM5 | RAC2 | | | |
| | | | RAC2 | | | | | | GRM7 | | | | |
| | | | RORC | | | | | | LRRTM2 | | | | |
| | | | STRA6 | | | | | | NRXN2 | | | | |
| | | | TBX5 | | | | | | PIK3R5 | | | | |
| | | | | | | | | | RASGRP1 | | | | |
| | | | | | | | | | SYT3 | | | | |
| | | | | | | | | | SYT8 | | | | |
| | | | | | | | | | UNC13A | | | | |
